# Host skin lipids trigger *MAT*-dependent mating, pathogenic hyphal growth, and parasexual reproduction of *Malassezia furfur*

**DOI:** 10.64898/2026.07.17.739196

**Authors:** Erica Patriarca, Bastien Tirtiaux, Márcia David-Palma, Anna Floyd Averette, Eduardo Gushiken-Ibañez, Yves Poumay, Salomé Leibundgut-Landmann, Joseph Heitman, Marco A. Coelho, Giuseppe Ianiri

## Abstract

The lipophilic yeast genus *Malassezia* dominates the human skin mycobiome and, while typically commensal, is associated with skin disorders in which hyphal cells are frequently observed. However, the mechanisms underlying hyphal differentiation and its contribution to pathogenesis remain poorly understood. Here, we show that hyphal growth in *Malassezia furfur* requires a complete mating-type system and is promoted by noncanonical reproductive interactions. Interspecific mating generates stable diploid hybrids with evidence of recombination, whereas intraspecific mating produces cell-fusion products that undergo recombination and haploidization, consistent with a parasexual cycle. Functional analyses reveal that the pheromone/receptor (*P/R*) locus governs cell recognition and fusion, while the homeodomain (*HD*) locus is required for hyphal development despite having lost canonical mating-type determination. Leveraging *in vitro* and *in vivo* skin models, we show that hyphal cells invade the epidermis more efficiently and elicit stronger inflammatory responses than yeast cells. Together, these findings link parasexual-like reproduction, mating-type loci function, and morphogenesis in *Malassezia* to pathogenic interactions with the host.

## Introduction

The skin microbiome of warm-blooded vertebrates comprises diverse microbial communities that interact with host tissues and contribute to both homeostasis and disease. Within this ecosystem, the fungal component (the mycobiome) is frequently dominated by the lipophilic basidiomycetous yeast genus *Malassezia*. In humans, *Malassezia* spp. are by far the most abundant fungal colonizers of the skin and thrive in sebum-rich areas such as the scalp, back, or neck. This association reflects a strong reliance on external lipids, consistent with the loss of fatty acid synthase genes across the genus ^1,2^. Among the 21 species formally described, *M. globosa* and *M. restricta* are the most commonly found on humans, followed by *M. sympodialis* and *M. furfur* ^2^. In addition to their role as commensals*, Malassezia* species can protect their hosts from *Staphylococcus aureus*, a bacterium implicated in atopic dermatitis pathogenesis, through protease-mediated biofilm degradation by *M. globosa*^3^, and secretion of a hydroxy palmitic acid isomer by *M. sympodialis* ^4^. Elevated *Malassezia* abundance on the skin also appears to limit colonization by the globally emerging multidrug-resistant fungal pathogen *C. auris* ^5^. Last, *M. furfur* can convert tryptophan into indole metabolites that activate the aryl hydrocarbon receptor (AhR), improving barrier function and controlling inflammation in diseased skin ^6^.

Despite these protective roles, *Malassezia spp*. are also implicated directly or indirectly in several skin pathologies, including atopic dermatitis, *Malassezia* folliculitis, seborrheic dermatitis (SD), and pityriasis versicolor (PV) ^7,8^. PV is characterized by *Malassezia* predominantly in the hyphal rather than the yeast form. While such morphology was thought to be PV-specific, hyphal structures have also been observed in SD skin lesions, indicating that hyphal growth is not restricted to PV ^9^. The ability of specific *M. furfur* strains to produce hyphae has recently been demonstrated in a reconstructed human epidermis (RHE) infection model, where hyphae invaded the cornified layer and induced strong inflammatory keratinocyte responses, while neither response was observed for yeast-form strains ^10,11^. Despite establishing strain-level differences in hyphal growth potential, and a key role of hyphae in epidermal invasion and pathogenicity, the genetic mechanisms governing the yeast-to-hyphae transition remain elusive.

In most basidiomycetes, the transition from yeast to hyphal growth is linked to sexual reproduction between two compatible mating partners, and it is controlled by two mating type (*MAT*) loci ^12^. The first is *MAT A*, which is generally biallelic and encodes pheromones (*MFA*) and pheromone receptors (*PRA*) that govern cell recognition and fusion. The second is *MAT B*, which encodes adjacent, divergently transcribed homeodomain genes (*bE*/*bW* or *HD1*/*HD2*), often multiallelic, that typically form a functional heterodimer only when bE and bW are derived from different *MAT B* alleles. This heterodimer acts after syngamy to drive developmental programs, such as hyphal development and host invasion, as demonstrated in the plant pathogen *Ustilago maydis* ^13–15^. Compatibility at both loci is thus typically required for sexual development ^12^. The *MAT A* and *MAT B* loci can be unlinked on distinct chromosomes (tetrapolar system), or linked on the same chromosome with suppression of recombination between the two loci (bipolar system). An intermediate pseudobipolar system, identified in some *Malassezia* species, retains both loci on the same chromosome but sufficiently far apart that recombination occurs ^16^.

Genome sequencing across *Malassezia* first revealed that some *M. furfur* strains have genomes approximately twice the size of other isolates, suggesting a non-haploid status and possible hybridization ^17^. A subsequent study identified two parental haploid lineages within *M. furfur* (P1 and P2) hypothesized to represent two distinct species, and two derived hybrid lineages (H1 and H2, diploid or aneuploid) resulting from hybridization of P1 and P2 strains ^18^. Genome analysis further revealed a pseudobipolar mating system with two *MAT A* and four *MAT B* alleles; P1 and P2 isolates are either *MAT a1* or *a2*, while *MAT B* is fixed within each lineage (*b2* in P1, and *b1 in* P2). H1 lineage strains carry *MAT a2b1b4* or *a2a2b1b4,* and are diploid or aneuploid, suggesting chromosome loss and/or loss of heterozygosity (LOH) during their evolution. Conversely, H2 lineage strains are diploid with a fully compatible set of *MAT* alleles, namely *MAT a1a2* and *b3b4* ^16,18^.

With the aim of linking *MAT* composition to hyphal formation in *M. furfur*, H2 lineage strains were grown in a number of conditions in earlier studies ^16,18^, but hyphae were never observed. The first evidence for *MAT* genes involvement in hyphal development came from a genetic engineering experiment in which insertion of compatible *MAT A* and *B* alleles (*a1b2*) or *MAT A* only (*a1*) into a haploid *M. furfur* strain (*MAT a2b1*) triggered robust hyphal formation under diverse growth conditions. The resulting strains were called “*A*on *B*on” (or “solo”; GI156) and “*A*on *B*off” (GI166) ^16^ following the nomenclature of *U. maydis* solopathogenic strains ^19^. Despite sharing the same *MAT* allelic composition as H2-lineage strains, only the engineered strains produced hyphae in culture. However, during a follow-up study assessing virulence in an RHE infection model, we noticed that, besides the “*A*on *B*on” and “*A*on *B*off” strains, some wild type *M. furfur* isolates also formed hyphae whereas others remained as yeast cells ^10^. Further experiments revealed that the hyphal growth in RHE was restricted to H2 lineage strains (eg CBS7019 and CBS6000), which are diploid and harbor a fully compatible set of *MAT* alleles; in contrast, H1 lineage strains and P1 or P2 haploid strains carrying an incomplete set of *MAT* alleles, failed to form hyphae ^10^. Together, these observations provided indirect evidence linking *MAT* composition to hyphal growth, but left unresolved whether sexual reproduction occurs in *Malassezia* and how it contributes to the yeast-to-hyphae transition.

Here, we define the nutritional conditions that induce hyphal growth in *M. furfur,* and show that they also promote cell-cell fusion. Mating between *MAT*-compatible strains from the two parental lineages generated diploid fusion products that acquired the ability to produce hyphae, whereas within-lineage crosses also yielded hypha-forming fusion products, although with variable ploidy that decreased to haploid or near haploid upon serial passaging, reminiscent of a parasexual-like cycle. Functional genetic analysis confirms a role for *MAT A* in cell recognition and fusion, as in other basidiomycetes, and for a single *MAT B* allele in hyphal development, without requiring a biparental HD protein combination. With *in vitro* and *in vivo* models, we demonstrate that *M. furfur* hyphae invade the skin more efficiently and elicit stronger host inflammatory responses than yeast cells, supporting a key role for hyphal growth in pathogenesis. Overall, our findings identify a noncanonical parasexual reproductive route in *Malassezia* that links cell fusion and genome plasticity with hyphal growth and pathogenic interactions with the host.

## RESULTS

### Olive oil promotes *Malassezia furfur* hyphal growth

Because some *M. furfur* strains undergo hyphal growth in the RHE infection model but not under laboratory conditions that instead readily induce hyphal growth in the engineered “*A*on” strains, we reasoned that the RHE model contains additional morphogenetic cues. We therefore dissected this system component by component with the hybrid strain CBS7019 as a representative isolate (**Supplementary Table 1**). Across combinations of individual RHE components and culture parameters, we consistently identified olive oil as the only factor associated with hyphal growth of CBS7019; olive oil was specifically included as an external lipid source to support *Malassezia* growth in the RHE model^10^. Remarkably, inoculating *M. furfur* CBS7019 into a drop of olive oil on a minimal medium (MM) agar plate at 30°C was sufficient to induce hyphal growth without any other RHE component (**Fig. 1a**). Hyphae were observed from the second day of incubation, and their abundance, length, and branching, increased over time (**Fig. 1b**). Individual hyphae showed a clearly distinguishable initial cell, septate hyphal compartments, and an apical growth cone (**Fig. 1c**). The presence of both a carbon source (glucose) and a nitrogen source (ammonium sulphate, or ammonium nitrate) was essential for hyphal development in the presence of olive oil, with growth being most robust when both nutrients were present in MM agar, a condition we hereafter refer to as filamentation medium (**Supplementary Table 2**).

**Figure 1.**
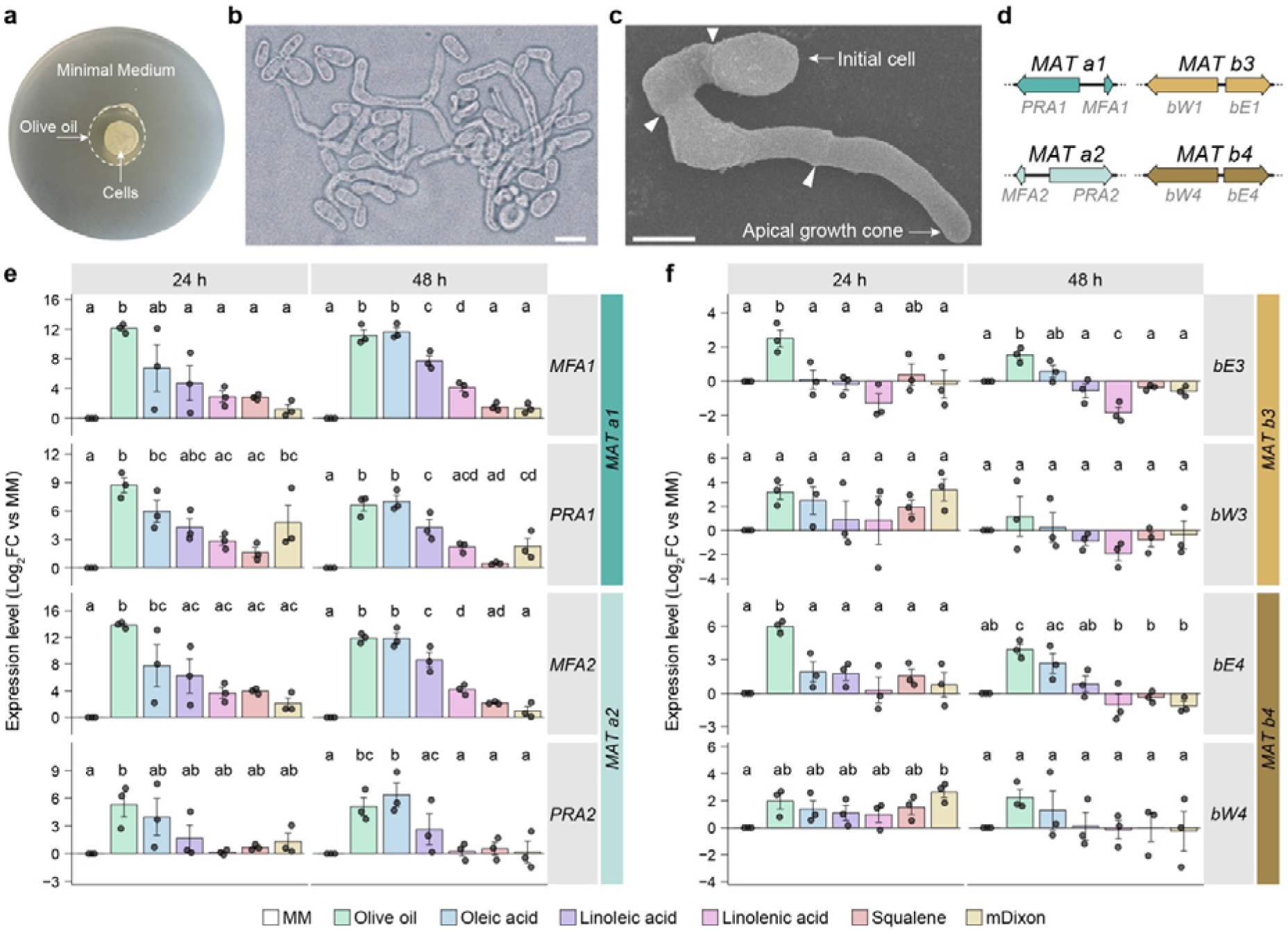
*M. furfur* hyphal formation and *MAT* gene expression under lipid-inducing conditions. **(a)** Experimental setup to induce hyphal growth in *M. furfur* CBS7019. Cells were resuspended in water and spotted within a drop of olive oil placed on filamentation medium (MM agar). **(b)** Hyphae of CBS7019 recovered after 3 days of incubation at 30°C (conditions as in panel **a**). **(c)** SEM image of a CBS7019 hypha; the initial cell, apical growth cone, and the septa (arrowheads) are indicated. **(d)** Schematic of the *MAT* locus organization in CBS7019, showing the two *MAT A* alleles (*a1* and *a2*) and two *MAT B* alleles (*b3* and *b4*) present in this diploid strain. Because the genome assembly of CBS7019 is not phased, the allelic combinations between *MAT A* and *MAT B* on each haplotype remain unknown. **(e, f)** RT-qPCR expression levels of **(e)** *MAT a1* and *a2* genes, and **(f)** *MAT b3* and *b4* genes in *M. furfur* CBS7019 after 24 h and 48 h on filamentation medium alone (MM; reference condition) or MM supplemented with olive oil, oleic acid, linoleic acid, linolenic acid, or squalene, or on mDixon. Expression levels are shown as log_2_ fold change (FC) relative to MM, calculated using the ΔΔCt method with *TUB2* as endogenous reference; MM log2 FC is defined as 0. Bars show the mean log_2_FC across biological replicates (n = 3); error bars indicate the standard error of the mean; dots indicate individual biological replicate values, each derived from the mean of three technical replicates. Statistical analysis was performed for each gene and time point on ΔCt values using a one-way ANOVA with condition as the main factor and biological replicate as a blocking factor, followed by Tukey-adjusted pairwise comparisons. Letters above bars indicate Tukey groupings (α = 0.05), with shared letters indicating no significant differences. Full pairwise comparison results are provided in Additional file 1.

Analysis of nuclear dynamics and hyphal architecture revealed that *M. furfur* CBS7019 diploid yeast cells are uninucleate, whereas hyphae are monokaryotic, with nuclei typically observed near the center of each hyphal compartment, and frequently near the septa (**Supplementary Fig. 1a-e**). Two nuclei were often observed near the terminal septum of growing hyphae (**Supplementary Fig. 1f-g**), suggesting that nuclear division occurs within a hyphal compartment before nuclear migration into the adjacent compartment. Consistent with a monokaryotic organization, true clamp connections (hook-like structures typical of basidiomycetes that ensure nuclear distribution in dikaryotic hyphae) were not observed in CBS7019 or, upon re-examination, in the self-filamenting strains GI156 and GI166 (**Supplementary Fig. 2**), although hook-like septal protrusions were occasionally noted in these strains^16^. Together, these observations establish olive oil as a cue for monokaryotic, clamp-less hyphal growth in *M. furfur*.

### Unsaturated fatty acids promote *MAT* gene expression under hyphae-inducing conditions

To define the lipid component(s) that triggers hyphal growth, we grew *M. furfur* CBS7019 on filamentation medium supplemented with a panel of commercial oils. In addition to olive oil, other oils induced hyphal formation, including jojoba, peanut, corn, sunflower, colza, and stearic oil (a commercial preparation rich in unsaturated fatty acids) while others, such as coconut oil, did not (**Supplementary Table 3**). A common feature between oils that promoted hyphal formation is their abundance in unsaturated fatty acids. Supplementation with individual unsaturated fatty acids (oleic, linoleic, linolenic acids) and the polyunsaturated hydrocarbon squalene, induced hyphal formation in CBS7019, but only observed after prolonged incubation. The addition of Tween 60 (1%) to support better *M. furfur* grow in the presence of individual unsaturated fatty acids or squalene, enhanced hyphal formation, although less strongly than olive oil. By contrast, neither Tween alone nor the saturated fatty acid palmitic acid induced hyphal formation (**Supplementary Table 3**).

The *M. furfur* H2-lineage strains CBS7019 and CBS6000, both diploid and capable of hyphal growth, carry a fully compatible *MAT* configuration, specifically *MAT a1a2, and b3b4* ^16,18^ (**Fig. 1d**), which we hypothesized might contribute to hyphal growth. To test this, we first assessed *MAT* gene expression by RT-qPCR in CBS7019 grown on filamentation medium supplemented with olive oil, oleic acid, linoleic acid, linolenic acid, or squalene, or grown in mDixon. At 24 h, olive oil induced the highest expression of *MAT A* genes, with significantly increased expression of *MFA1*, *PRA1*, and *MFA2* relative to the MM reference and most other conditions. Oleic acid also induced high *MAT A* gene expression, although responses were more variable and several pairwise comparisons were not statistically significant (**Fig. 1e**). Conversely, linoleic acid, linolenic acid, and squalene showed modest *MAT A* gene induction and were not statistically distinguishable from MM at 24 h (**Fig. 1e**). After 48 h, olive oil and oleic acid still induced the highest *MAT A* gene expression, while linoleic acid showed intermediate levels; linolenic acid, squalene, and mDixon, showed variable and generally modest induction, frequently indistinguishable from MM or from each other (**Fig. 1e**). For the *MAT B* locus*, bE3* and *bE4* alleles showed their strongest and most consistent upregulation with olive oil at 24 h and 48 h, and with oleic acid 48 h, whereas *bW3* and *bW4* did not differ significantly across conditions (**Fig. 1e**).

These findings show that unsaturated fatty acids promote both hyphal formation and *MAT* gene expression in *M. furfur*, with olive oil eliciting the strongest and most consistent response, possibly reflecting a synergistic contribution of multiple lipid components. Based on these findings, all of the subsequent experiments were carried out on filamentation medium with cells placed in a drop of olive oil, unless stated otherwise.

### Compatibility at *MAT A* is required for *M. furfur* hyphal growth

To test whether the *MAT* locus controls the yeast-to-hyphae transition, we first screened a panel of *M. furfur* strains spanning different *MAT* genotypes and lineages (**Supplementary Table 1**) for hyphal formation with the olive-oil droplet assay (as in **Fig. 1a**), scoring cell morphology by light microscopy. The panel included haploid P1 and P2 lineage strains, H1-lineage hybrids (CBS1878 and CBS4171) and H2-lineage hybrids (CBS7019 and CBS6000; positive controls), the self-filamenting strains GI156 and GI166, and the newly engineered strain #216, generated by inserting the transgene *MATa1-NEO-MATb4* into the P2 haploid strain PM315 (*MAT a1b1*). Robust hyphae formation was restricted to strains carrying compatible *MAT A* alleles (CBS7019, CBS6000, GI156, and GI166), whereas those carrying an incomplete *MAT* system (haploid strains and H1-lineage hybrids)^18^ remained in the yeast form. Occasional pseudohyphal-like cells were observed in haploid strains (**Supplementary Fig. 3**), but were readily distinguishable from true hyphae, which were multicellular, with septa, and generally longer and wider. Together, these results are consistent with a requirement for *MAT A* compatibility for hyphal growth.

Next, we performed mating assays under the same conditions with four *M. furfur* haploid strains (PM315, CBS9574, CBS14141, and CD866; **Supplementary Table 1**), representing both P1 and P2 lineages and collectively covering both *MAT A* and *B* alleles, allowing interspecific P1 × P2 (CBS14141 × CBS9574 and CD866 × PM315) and intraspecific crosses (P1 × P1: CD866 × CBS9574; P2 × P2: CBS14141 × PM315). Although hyphae were observed in several conditions, results were variable across replicates and timepoints; we therefore developed a selection-based assay to directly recover and quantify cell fusion products.

### *MAT A-*compatible *M. furfur* strains undergo cell fusion generating fusion products that gain the ability to produce hyphae

Hyphal growth appeared to be the result of cell fusion events controlled by the *MAT* loci. To directly test this, we utilized *Agrobacterium*-mediated transformation to generate haploid *M. furfur* strains PM315, CBS9574, CBS14141, and CD866 carrying dominant selectable markers conferring resistance to nourseothricin (NAT) or G418 (geneticin; NEO). One stable NAT- or NEO-resistant transformant was selected for each strain, with the exception of PM315, for which two independent NEO-resistant transformants were analyzed (**Supplementary Table 1**). Marked strain pairs were co-inoculated in the olive-oil droplet assay (**Fig. 2a**) and after 3 days of incubation at 30°C, the co-cultures were plated onto mDixon agar containing both NAT and NEO. Double drug-resistant colonies were recovered only from crosses involving parental strains compatible at *MAT A* (with or without *MAT B* compatibility), whereas crossing with compatibility only at *MAT B* did not yield double drug-resistant colonies (**Fig. 2b**). Across five independent positive assays, double drug-resistant colony counts were broadly consistent within each cross but differed among crosses, with crosses CBS14141-NAT#1 × CBS9574-NEO#1 (P2 × P1) and CD866-NAT#1 × CBS9574-NEO#1 (P1 × P1) yielding significantly more putative fusion products (**Fig. 2c**; P <0,0001). Further experiments indicate that cell fusion can occur within 24 h (**Supplementary Fig. 4**). Cell fusion was also tested on filamenting medium supplemented with the same individual lipids as in figure 1e-f; besides olive oil (positive control), oleic acid yielded the highest recovery of double-drug resistant colonies, while linoleic and linolenic acid yielded only 1 and 4 colonies, respectively, in a single cross (**Supplementary Table 4**).

**Fig. 2.**
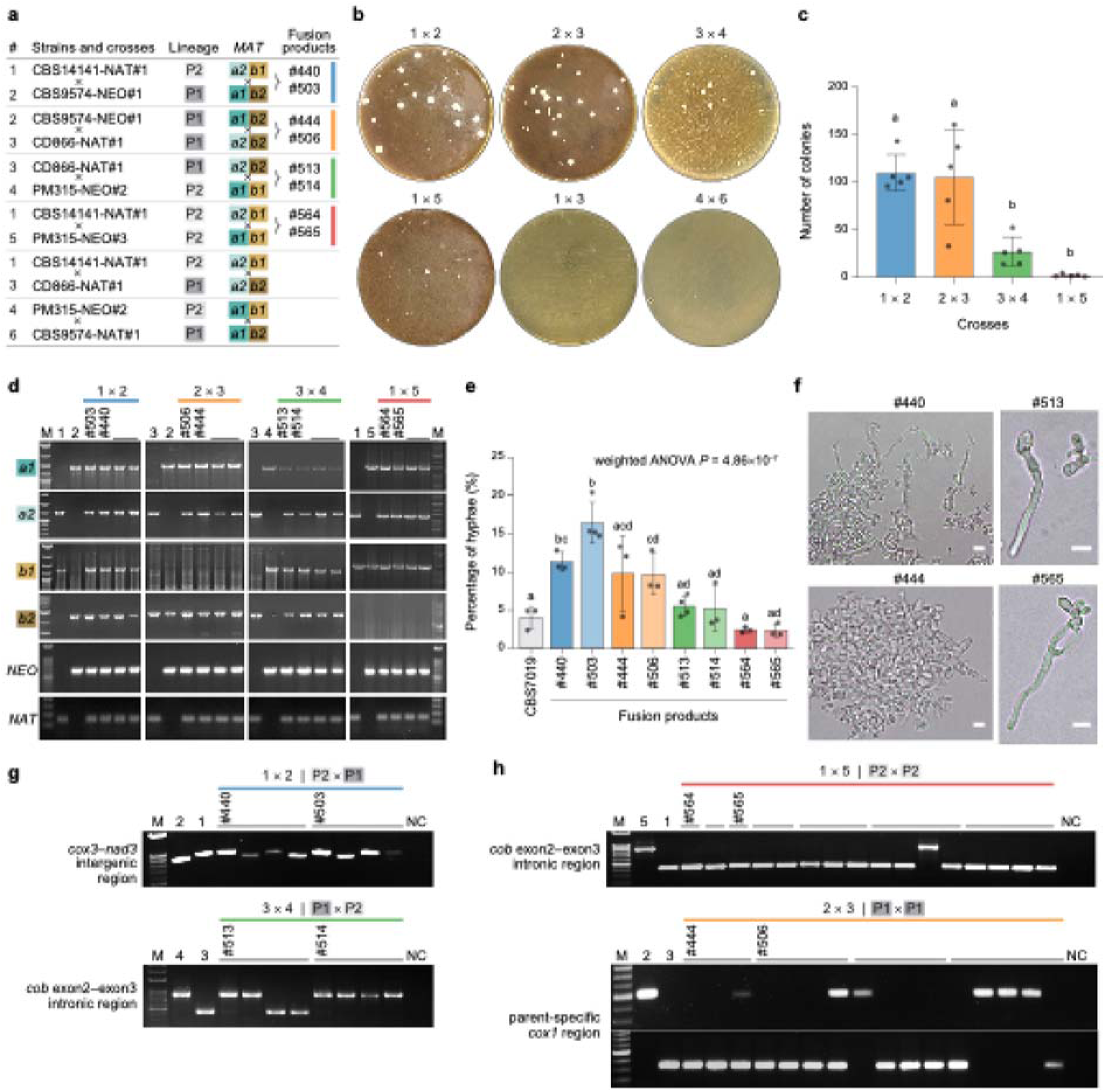
*MAT A-*compatible *M. furfur* strains undergo cell fusion and generate fusion products that gain the ability to form hyphae. **(a)** Summary of the haploid parental strains, lineage assignment (P1 or P2), *MAT* allele genotypes, and the fusion products recovered from each cross (coded as 1×2, 2×3, 3×4, and 1×5, matching the parental numbering shown in the table). **(b)** Representative double-drug selection plates from the indicated inter-specific (P1 × P2) and intra-specific (P1 × P1 or P2 × P2) crosses showing recovery of fusion-product colonies. In cross 1×5, only one colony was recovered in the plate shown. **(c)** Quantification of colony formation under double-drug selection across independent crosses. Bars show the mean and error bars indicate SD; points represent individual plates from five independent mating assays per cross, each plated in duplicate or triplicate. Differences across crosses were determined by one-way ANOVA followed by Tukey’s multiple-comparison test. Full pairwise comparison results are provided in Additional file 2. **(d)** PCR genotyping of the parental haploid strains and selected fusion products to assess the *MAT* genotype (*a1*, *a2*, *b1*, *b2*) and the *NAT/NEO* drug-resistance markers. Fusion products selected for downstream analyses are labeled (#). **(e)** Filamentation rate after 72 h of incubation on filamentation medium with olive oil, reported as the percentage of hyphal cells, for the natural hybrid strain CBS7019 and selected fusion products. Bars show the mean across independent measurements and error bars indicate SD; points represent individual measurements (*n* = 3 per strain, except #503 and #513 where *n* = 4). Statistical differences across groups were assessed by weighted one-way ANOVA (weights = total cells scored per measurement) followed by Tukey’s multiple-comparison test. Letters above bars indicate Tukey groupings (α = 0.05), with shared letters indicating no significant differences. Full pairwise comparison results are provided in Additional file 3. **(f)** Representative microscopy images of filamentation phenotypes for fusion products from interspecific (#440 and #513) and intraspecific (#444 and #565) crosses; images of other fusion products are available in **Supplementary Fig. 5.** Scale bars, 5 μm. **(g–h)** Mitochondrial inheritance in representative fusion products assessed by PCR with markers distinguishing parental mitochondrial genotypes; amplicon regions are labeled to the left of each gel. Parental strains are included as reference; NC, no-template control; M, ladder.

Because double selection confirms acquisition of *NAT* and *NEO* but does not resolve *MAT* allele composition, we PCR-genotyped four randomly-selected double drug resistant colonies from each cross. Colonies from crosses with strains compatible at both *MATA* and *MATB* loci were PCR positive for the expected *MAT* alleles (*a1, a2, b1, b2*) as well as the selectable markers *NAT* and *NEO* (**Fig. 2d**). Likewise, colonies from crosses with strains compatible at *MAT A* but not *MAT B* were PCR-positive for *MAT a1, a2, NAT,* and *NEO*, and carried the same *MAT B* allele as the parental strains (*b1* for CBS14141-NAT#1 × PM315-NEO#3; and *b2* for CD866-NAT#1 × CBS9574-NEO#1) (**Fig. 2d**). Regardless of *MAT B* genotype, all double drug-resistant colonies PCR-positive for both *MAT a1* and *a2* formed hyphae (**Fig. 2e, f**).

Quantification of hyphal formation for two independent fusion products from each cross showed that fusion products #440 and #503 (P1 × P2) and #444 and #506 (P1 × P1) had the highest levels (≥10%), followed by #513 and #514 (P1 × P2). The natural hybrid CBS7019 displayed about 4% hyphal formation, whereas fusion products #564 and #565 (P2 × P2) showed about 2%, the lowest levels recorded (**Fig 2e**). These values likely represent conservative estimates of hyphal formation, as tangled hyphae could not be reliably counted and washing steps may have perturbed the samples. Microscopic analysis of a representative fusion product from each cross revealed some variation in hyphal morphology but no consistent traits distinguishing intra- from inter-specific fusion products (**Fig 2f; Supplementary Fig. 5**).

Mitochondrial inheritance in *M. furfur* fusion products was assessed with previously reported mitogenome polymorphic markers ^20^. Strains selected for this analysis were confirmed as cell fusion products by PCR genotyping of *MAT* allele markers, and by their ability to form hyphae. In the interspecific cross CBS14141-NAT#1 (P2, *a2b1*) × CBS9574-NEO#1 (P1, *a1b2*), primers targeting the *cox3–nad3* intergenic region showed that 3/8 fusion products carried the CBS9574 mitogenome and 5/8 carried the CBS14141 mitogenome (**Fig. 2g**). In the interspecific cross CD866-NAT#1 (P1, *a2b2*) × PM315-NEO#2 (P2, *a1b1*), primers targeting the *cob* exon2–exon3 intronic region indicated that 6/8 fusion products carried the PM315 mitogenome and 2/8 carried the CD866 mitogenome (**Fig. 2g**). In the intraspecific cross CBS14141-NAT#1 (P2, *a2b1*) × PM315-NEO#3 (P2, *a1b1*), 15/16 fusion products carried the CBS14141 mitogenome (**Fig. 2h**). For the intraspecific cross CD866-NAT#1 (P1, *a2b2*) × CBS9574-NEO#1 (P1, *a1b2*), existing markers could not discriminate parental mitochondria; we therefore designed parental-specific primers for *cox1* (**Supplementary Fig. 6**), which showed that 10/16 fusion products carried the CD866 mitogenome, 4/16 carried the CBS9574 mitogenome, and 2/16 were positive for both, potentially reflecting residual heteroplasmy or mixed template amplification (**Fig. 2h**). Overall, these results show that mitochondrial inheritance in P1 × P2 interspecific crosses was variable and consistent with biparental inheritance, whereas in intraspecific crosses showed a bias toward inheritance from the *MATa2* parent, similar to uniparental inheritance reported in taxonomically related fungi ^21^.

To visualize cell fusion events directly, an mCherry-tagged derivative of the *M. furfur* strain CBS14141 was generated by *Agrobacterium*-mediated transformation and crossed with wild type strains CBS9574 (P2 × P1) or PM315 (P2 × P2). Microscopic examination of CBS14141-mCherry × CBS9574 mating cultures revealed conjugation tubes connecting mCherry-tagged and untagged cells (**Supplementary Fig. 7a**). In the intraspecific cross, we captured a conjugation tube emerging from a pseudohyphal cell of PM315 and connecting to a yeast cell of CBS14141-mCherry (**Supplementary Fig. 7b**). Cell-cell connections were frequently observed involving pseudohyphae (**Supplementary Fig. 7c**), but were also seen between yeast cells (**Supplementary Fig. 7d**). Conjugation tubes appeared morphologically distinct from pseudohyphae, suggesting they might represent a specialized mating structure. Finally, double drug resistant putative products generated by crossing CBS14141-mCherry with CBS9574-NEO#1 (strain #524) or PM315-NEO#3 (strain #644), displayed robust hyphal formation and cytoplasmic mCherry fluorescence (**Supplementary Fig. 7e-f**), consistent with m-Cherry-positive cell fusion products giving rise to hyphal growth.

### The fusion products from interspecific crosses are diploid hybrids

To determine the ploidy and genome composition of fusion products from the two interspecific crosses, we analyzed the parental strains and two fusion products from each cross by flow cytometry (FACS) and Illumina sequencing (**Fig. 3a**). Mapping Illumina reads to a parental reference genome for each cross revealed chromosome-wide heterozygosity (**Fig. 3b-c**), except for small LOH tracts in fusion product #440 on chr. 2 (∼9.2 kb) and in #503 on chr. 6 (∼5.6 kb); a larger LOH region (∼152 kb) was also detected at the left end of chr. 5 in #503. Consistent with their hybrid origin, FACS confirmed that all four fusion products were diploid whereas the parental strains were haploid, as expected (**Fig. 3d**). De novo genome assemblies further supported this, revealing genome sizes exceeding 16 Mb (**Supplementary Table 5**) in all four fusion products, approximately twice the size of the haploid parental genomes (∼8 Mb). BLAST searches confirmed the presence of parental *MAT* allele markers (*a1*, *a2*, *b1*, and *b2*), and the *NAT* and *NEO* cassettes. The loci disrupted by random insertion of *NAT* and *NEO* in the parental strains are reported in **Supplementary Fig. 8**; although these insertions may influence cell fusion frequency or hyphal formation levels, they do not appear to impair either process. Together, these results confirm that the fusion products from interspecific crosses are diploid hybrids (allodiploids), comparable in genome size to naturally occurring *M. furfur* hybrids but with markedly lower levels of LOH ^18^.

**Fig. 3.**
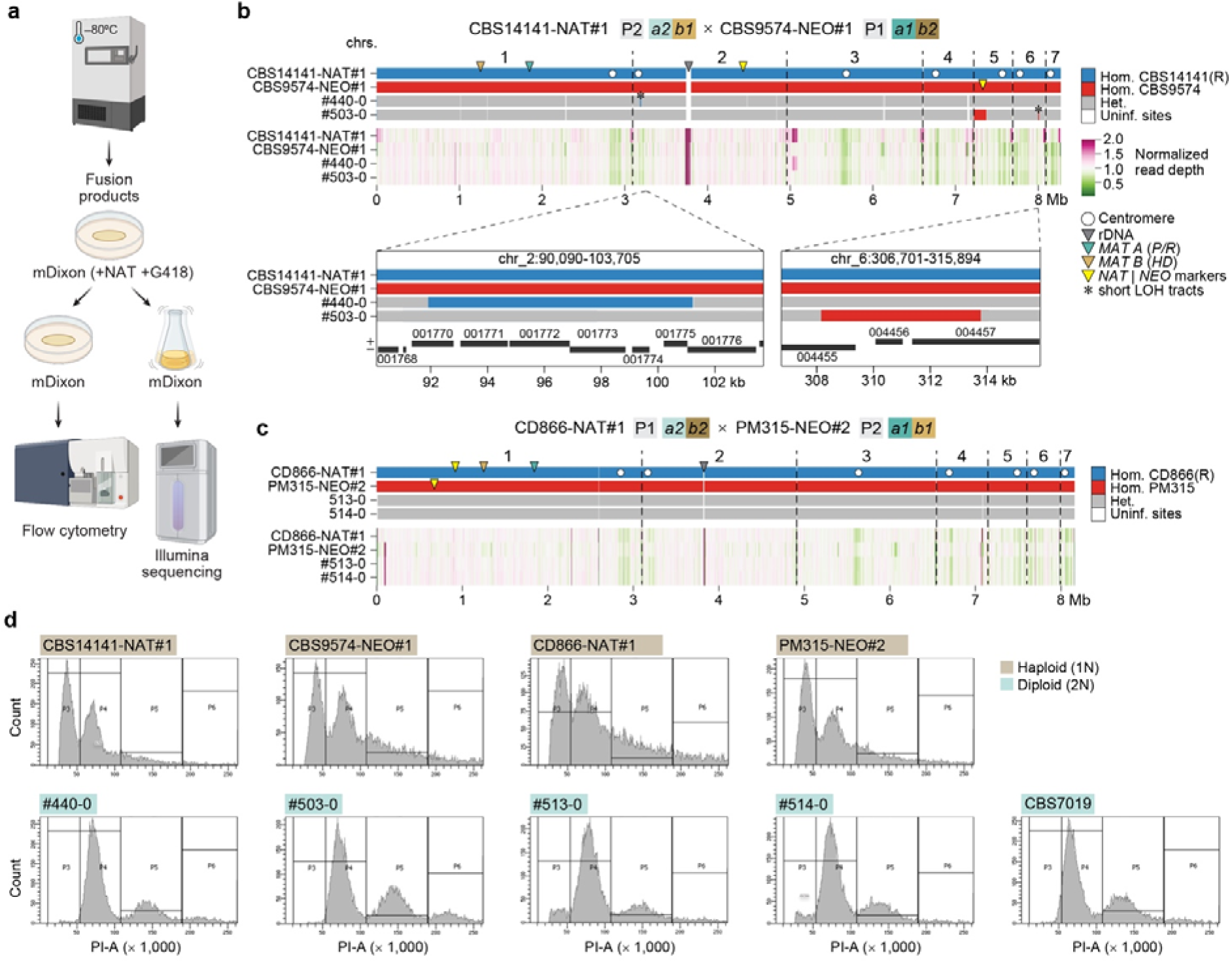
Genome composition and ploidy of *M. furfur* fusion products from interspecific P1 × P2 crosses between marked, *MAT*-compatible strains. **(a)** Workflow for characterization of fusion products by flow cytometry and Illumina sequencing. Fusion products were revived from frozen stocks on mDixon agar supplemented with NAT and G418, then grown in liquid mDixon for DNA extraction, or on solid mDixon for flow cytometry. **(b, c)** Genotype (top tracks) and normalized read-depth (bottom tracks) profiles for two independent fusion products from each interspecific cross: **(b)** CBS14141-NAT#1 × CBS9574-NEO#1 (#440 and #503) and **(c)** CD866-NAT#1 × PM315-NEO#2 (#513 and #514). Genotypes were inferred from parent-informative SNPs mapped to the indicated parental reference genome (R) and classified as homozygous reference (blue), homozygous alternate parent (red), heterozygous (gray), or uninformative (i.e. regions identical between both parental); adjacent variants of the same genotype class separated by ≤ 5 kb were merged. Read depth was calculated in non-overlapping 5-kb windows and normalized to the genome-wide median. Arrows indicate the reference-coordinate positions of the rDNA locus, T-DNA insertions, and *MAT A* and *MAT B* loci, as indicated in the key; T-DNA insertion sites were inferred from strain-specific *de novo* assemblies and projected onto the reference genome. White circles depict centromeres, and dashed lines indicate chromosome boundaries. Small regions of read-depth variation may reflect copy-number differences or parental divergence affecting read mapping. Inserts in **b** show localized LOH tracts with gene models lifted from the *M. furfur* CBS14141 annotation (+ and – indicate strand orientation). **(d)** Flow cytometry profiles of parental strains, fusion products, and the diploid control CBS7019. Haploid and diploid profiles are labeled in brown and in light green, respectively.

### Intraspecific fusion products undergo ploidy reduction and recombination upon passaging

Unlike the diploid interspecific fusion products, the intraspecific fusion products #444 and #506 (P1 × P1) and #564 and #565 (P2 × P2) initially appeared haploid by FACS yet retained extensive genome-wide heterozygosity characteristic of diploid hybrid genomes (**samples 0a; Supplementary Fig. 9)**. This discrepancy prompted us to investigate whether ploidy reduction had occurred during sample handling, specifically, during passaging in non-selective medium. To test this, we performed FACS and Illumina sequencing on two sample sets: (i) the same fusion products revived from frozen stocks on selective mDixon medium (**Fig. 4**; samples 0b), and (ii) three single colonies derived from each fusion product, isolated and passaged three times on non-selective mDixon medium (**Figure 4a**; samples 1-3). These analyses showed that the intraspecific fusion products were indeed diploid or of mixed 1n/2n ploidy at the time of freezing; retrospective comparison with the original screening samples (**Supplementary Fig. 9,** samples 0a) further suggested that growth in non-selective conditions had already promoted LOH accumulation prior to the initial analysis (**Fig. 4; Supplementary Fig. 9**).

**Fig. 4.**
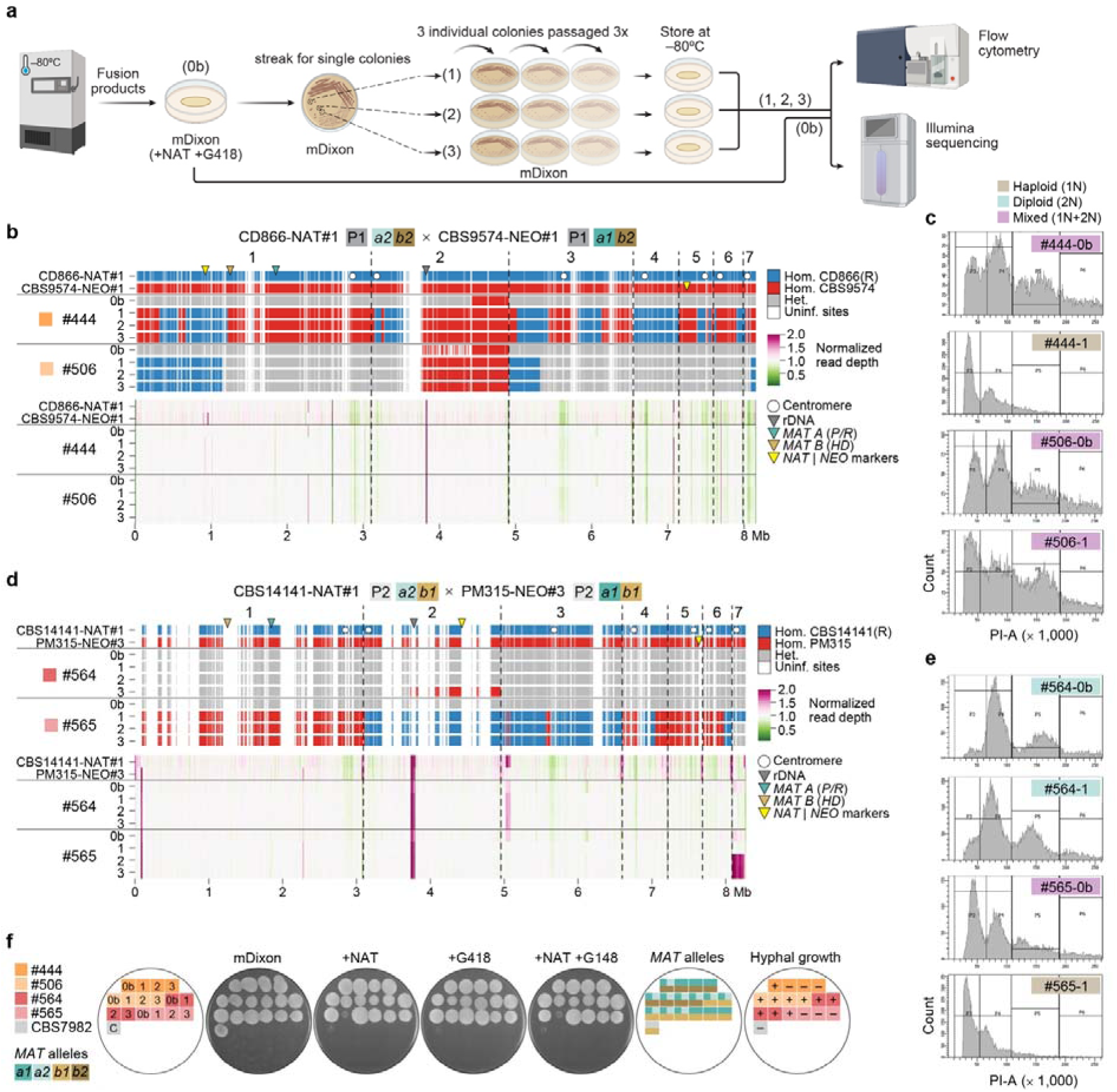
Ploidy reduction and recombination in *M. furfur* fusion products from intraspecific crosses between marked strains carrying different *MAT A* alleles but matching *MAT B* alleles. **(a)** Workflow for characterization of intraspecific fusion products by flow cytometry and Illumina sequencing. Revived fusion products (labeled as 0b) on double-drug (NAT+G418) mDixon plates were sampled directly for DNA extraction and flow cytometry. A portion of the same growth patch was streaked on non-selective mDixon, and three single-colony derivatives (labeled 1, 2, and 3) were passaged three times without selection before analysis by flow cytometry and Illumina sequencing. **(b–e)** Genome composition, normalized read-depth and flow cytometry profiles for the indicated intraspecific P1 × P1 and P2 × P2 fusion products and their passaged derivatives; genome tracks are shown in **b** and **d**, and representative flow cytometry profiles in **c** and **e**. Flow cytometry profiles of all strains are in **Supplementary Fig. 10**. Genotype classes and read-depth normalization are as described in Fig. 3; additional annotations are indicated in panel keys. **(f)** Drug resistance, *MAT* allele content, and hyphal growth of revived fusion products (0b) and passaged derivatives (1–3), spotted on mDixon with no drug, NAT, G418 or both drugs, and incubated at 30°C for 5 days. Hyphal growth was assessed on filamentation medium with olive oil and scored as present (+) or absent (–). *MAT* alleles and *NAT*/*NEO* markers were identified from *de novo* genome assemblies.

We then examined each cross in detail. For the P1 × P1 cross, fusion products #444 and #506 (samples 0b) showed extensive heterozygosity with a few LOH tracts, and mixed 1n/2n ploidy (**Fig. 4 b, c**). Passaged isolates derived from #444 were haploid and with recombinant genomes, having inherited markers from both parents across multiple chromosomes, with the exception of chromosome 4, which was inherited from a single parent. No differences were observed among the three #444-derived passaged isolates (**Fig. 4b-c; Supplementary Fig. 10**). In contrast, passaged isolates derived from fusion product #506 showed similar ploidy as the initial #506 fusion product, but accumulated additional LOH on chromosomes 1, 2, 3 and 7; notably, these LOH tracts overlapped with those observed in the initial #506 fusion product (sample 0a) growth in non-selective liquid medium, suggesting they may represent preferential sites of LOH accumulation (**Fig. 4b-c; Supplementary Fig. 9**).

The P2 × P2 fusion products showed a similar but not identical pattern. Fusion product #564 (sample 0b) and its passaged isolates were consistently heterozygous diploids, with the exception of isolate #564-3, which underwent LOH on a segment of chr. 2 encompassing the *NAT* marker, resulting in the loss of NAT resistance (**Fig. 4d**). For fusion product #565, the sample 0b was a heterozygous with mixed 1n/2n ploidy, while passaged isolates were haploid with recombinant genomes, showing recombination on chrs. 3, 4 and 6; chrs. 1 and 5 were inherited from one parent, chr. 2 from the other, and isolates #565-2 and #565-3 retained disomy of chr. 7 with persistent heterozygosity (**Fig. 4d-e**).

De novo genome assemblies of the passaged haploid isolates confirmed haploid genomes size (∼8 Mb), with the exception of isolates #565-2 and #565-3, which had slightly larger (∼8.34 Mb) and more fragmented assemblies, likely due to retention of chr. 7 from both parents (**Fig. 4d**). BLAST searches confirmed that all haploid isolates carried a single parental *MAT* configuration, rendering them unable to form hyphae, and had inherited both *NAT* and *NEO* resistance markers, consistent with independent segregation of the chromosomes carrying each marker during ploidy reduction (**Fig. 4f**); the only exception was isolate #564-3, which lost *NAT* resistance following LOH at the *NAT* locus as described above (**Fig. 4d**). Together, these findings demonstrate that intraspecific fusion products can undergo spontaneous ploidy reduction and recombination, generating recombinant haploid progeny, hallmarks of a parasexual cycle.

### Functional dissection reveals distinct *MAT A* and *MAT B* roles in cell fusion and hyphal growth

To define the specific contributions of the *MAT A* and *MAT B* loci to cell fusion and hyphal growth in *M. furfur,* we generated targeted deletions of the *MAT a2* (*MFA2*/*PRA2*) and *MAT b1* (*bE1/bW1*) loci in the wild-type CBS14141 background, and validated mutant genotypes by diagnostic PCR (**Fig. 5 a-b**). We then performed unilateral crosses (i.e., crosses where only one parent carries the deletion allele) between the CBS14141 mutants and marked parental tester strains to test whether *MAT A* or *MAT B* is required for cell fusion. Unilateral interspecific and intraspecific crosses of CBS14141 *a2*Δ (*mfa2*Δ/*pra2*Δ) with CBS9574-NEO#1 (*a1b2*) and with PM315-NEO#3 (*a1b1*), respectively, did not yield colonies on double-drug selective medium, indicating that *MAT A* is required for cell fusion (**Fig. 5c**). Conversely, unilateral crosses of CBS14141 *b1*Δ (*bW1*Δ/*bE1*Δ) with CBS9574-NEO#1 and with PM315-NEO#3 yielded double drug-resistant colonies confirmed by PCR to carry the expected *MAT* alleles (*a1*, *a2* and *b2* for the CBS14141 *b1*Δ × CBS9574-NEO#1 fusion products, and *a1*, *a2* and *b1* for the CBS14141 *b1*Δ × PM315-NEO#3 fusion products) and the *NAT* and *NEO* markers (**Fig. 5d**). Morphological analysis on filamentation medium revealed normal hyphal growth in fusion products recovered from both interspecific (strain #592) and intraspecific (strain #596) unilateral crosses (**Fig. 5e**), indicating that *MAT B* compatibility between mating partners is dispensable for both cell fusion and hyphal growth.

**Fig. 5.**
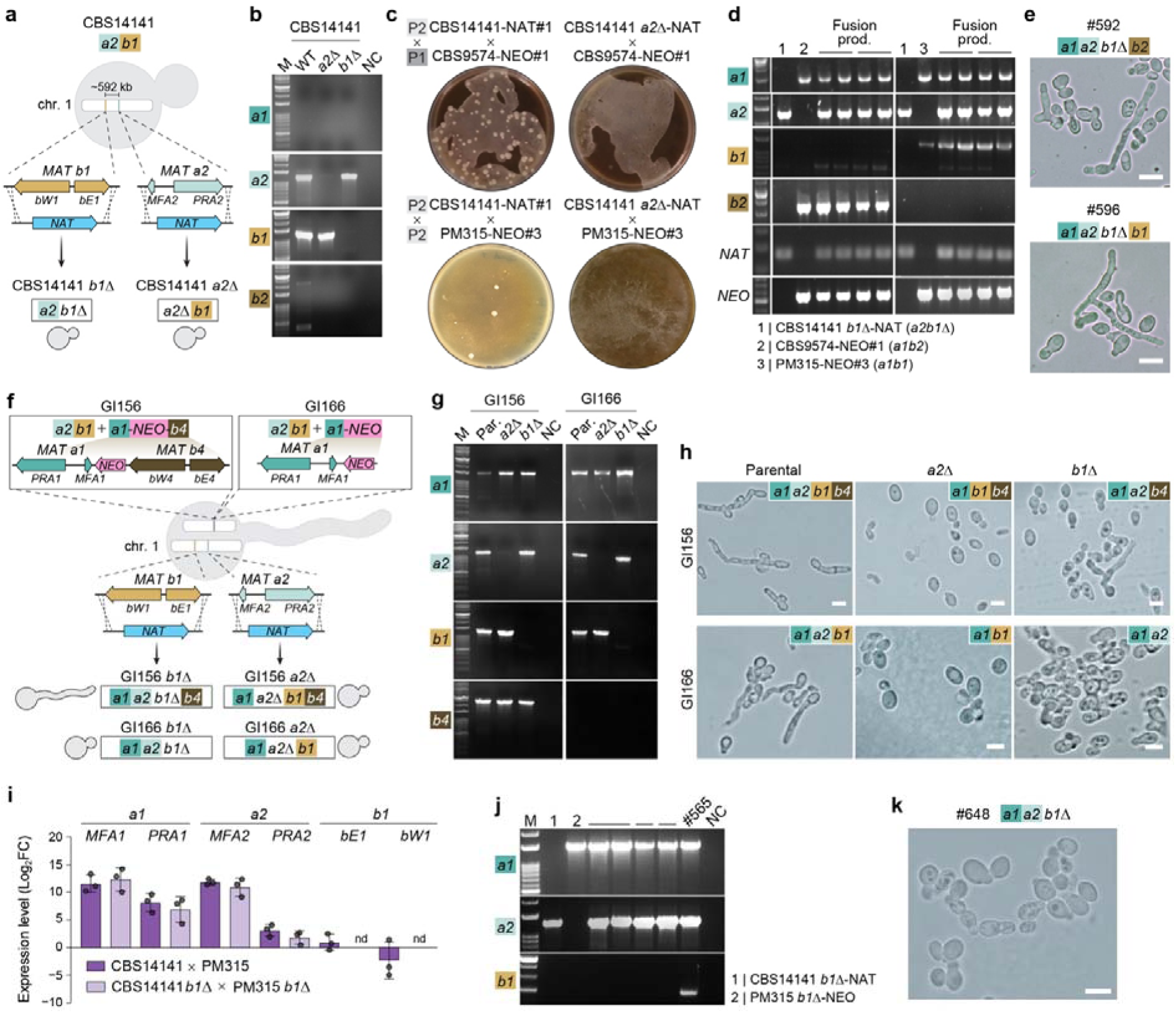
Functional role of the *MAT* A and *MAT B* loci in cell fusion and hyphal formation. **(a)** Schematic representation of *M. furfur* CBS14141 *mat a2*Δ and *mat b1*Δ mutants, generated by NAT-marker disruption of *MFA2/PRA2* (*MAT a2*) or *bW1*/*bE1* (*MAT b1*) on chr. 1; the resulting genotypes and yeast-form morphology, as assessed on filamentation medium, are shown. **(b)** PCR validation of *mat a2*Δ and *mat b1*Δ mutants to assess presence/absence of *MAT a1*, *a2*, *b1* and *b2* alleles; NC, no-template control; M, ladder. **(c)** Representative double-drug selection plates showing recovery of fusion products in the control crosses CBS14141-NAT#1 x CBS9574-NEO#1 (P2 × P1) and CBS14141-NAT#1 × PM315-NEO#3 (P2 × P2), and their absence in the corresponding crosses carried out with the CBS14141 *mat a2*Δ mutant. **(d)** PCR genotyping of fusion products from the unilateral crosses CBS14141 *mat b1*Δ × CBS9574-NEO#1 (P1 × P2), and CBS14141 *mat b1*Δ × PM315-NEO#3 (P2 × P2). Parental strains are numbered 1 - 3, with genotypes reported below the gels; black lines above the gels indicates fusion products from independent crosses. The weak band in the *MAT b1* panel is a non-specific amplicon. **(e)** Fusion products #592 and #596, obtained from unilateral crosses in **d**, from hyphae on filamentation medium with olive oil; their *MAT* configurations are indicated. **(f)** Schematic representation of the self-filamenting *M. furfur* strains GI156 and GI166 and their derived *mat a2*Δ and *mat b1*Δ mutants. Top: original constructs (*MAT a1-NEO-b4*, and *MAT a1-NEO*) inserted into strain CBS14141 to generate strains GI156 and GI166, respectively (Coelho et al., 2023); bottom: native *MAT a2* and *MAT b1* alleles on chr. 1, and their mutation achieved through homologous recombination using the *NAT* marker. The resulting mutants and their morphology, as assessed on filamentation medium with olive oil, are shown. **(g)** PCR validation of GI156 and GI166 and derived *mat a2*Δ and *mat b1*Δ mutants to assess presence/absence of the *MAT a1*, *a2*, *b1* and *b4* alleles; NC, no-template control; M, ladder. **(h)** Morphology of GI156 and GI166 and derived *mat a2*Δ and *mat b1*Δ mutants on filamenting medium with olive oil. **(i)** RT-qPCR expression levels of the genes *MFA1, PRA1, MFA2*, *PRA2*, *bE1* and *bW1* during the WT cross (CBS14141-NAT#1 × PM315-NEO#3) and the bilateral cross of derived *mat-b1*Δ mutants, both after 24 h on filamentation medium with olive oil, relative to the WT crosses on filamenting medium without olive oil (MM), used as the comparative condition. Ct values were normalized to the endogenous reference *TUB2* and converted to log2 fold change (log_2_FC) relative to MM using the ΔΔCt method; as MM is the calibrator, its log2 FC is defined as 0. Bars show mean log_2_FC across biological replicates (n = 3), with error bars indicating SEM; dots indicate individual biological replicate values, each the mean of three technical replicates. Statistical analysis was performed separately for each gene using an ANOVA model (condition as the main factor and biological replicate as a blocking factor), followed by Sidak pairwise comparisons. Nd, ‘not detected’. **(j)** PCR genotyping of fusion products from the bilateral cross CBS14141 *mat b1*Δ × PM315 *mat-b1*Δ; parental strains are indicated with numbers 1-2, with their genotypes shown next to the image; strain #565 served as positive control for *MAT B1*. Black line above the gel group fusion products from independent crosses. NC, no-template control; M, ladder. **(k)** Representative image of one fusion product (strain #648) from the bilateral cross in **j** on filamentation medium with olive oil. In all microscopic panels, the scale bar indicates 5 μm.

Next, to assess the contribution of each *MAT* locus to hyphal growth independently of cell fusion, we introduced the same *MAT a2::NAT* and *MAT b1::NAT* deletion cassettes into the self-filamenting strains GI156 (“*A*on *B*on”) and GI166 (“*A*on *B*off”) ^16^ (**Fig. 5 f-g**). In GI156, deletion of the native *MAT a2* locus abolished hyphal growth, whereas deletion of *MAT b1* (leaving *b4* intact) did not affect hyphal formation compared to the parental strain (**Fig. 5h**), indicating that *MAT A* compatibility is required for hyphal growth but *MAT B* allelic compatibility is dispensable. In GI166, deletion of *MAT a2* similarly abolished hyphal growth, while deletion of *MAT b1* loci — which in this background removes the only functional *MAT B* allele — also resulted in a yeast-only phenotype (**Fig. 5h**), demonstrating that at least one functional *MAT B* allele is required for hyphal growth independently of *MAT B* allelic compatibility. Gene expression analysis under hyphae-inducing conditions confirmed successful deletion of the target loci, with *MAT A* gene expression remaining strong and comparable across parental and mutant backgrounds; *MAT B* expression was overall low in all backgrounds, suggesting that its role in hyphal development does not depend on strong transcriptional induction under these conditions (**Supplementary Fig. 11**).

Given the yeast-only phenotype of the GI166 *b1*Δ mutant, we then tested whether *MAT B* is also required for hyphal growth in fusion products from genetic crosses. To test this, a *b1*Δ-*NEO* mutant was generated in PM315 (*a1b1*) and subjected to bilateral cross (i.e. in which both parents carry the deletion allele) with the CBS14141 *b1*Δ-*NAT* mutant. RT-qPCR analysis of mating cultures revealed high expression of *MFA1, PRA1, MFA2,* and *PRA2* with no significant differences relative to the wild type control cross, and the expected absence of *bE1* and *bW1* expression, confirming successful loss of *MAT b1* in both parents (**Fig. 5i**). PCR genotyping of recovered double drug-resistant colonies confirmed they were genuine fusion products, carrying *MAT a1* and *a2* alleles from each parent and lacking *MAT b1*, as expected from the bilateral deletion (**Fig. 5j**). Phenotypic analysis of a representative *b1*Δ fusion product (#648) revealed a yeast-only phenotype (**Fig. 5k**), demonstrating that *MAT B* is dispensable for cell fusion but required for hyphal formation. Taken together with the unilateral *b1*Δ cross results — where fusion products carrying a single *MAT B* allele (*b2* only in #592, *b1* only in #596) formed hyphae normally — these findings reveal that a single functional *MAT B* allele is sufficient to drive hyphal growth, regardless of allelic identity.

### Fusion products exhibit enhanced virulence-associated phenotypes relative to parental strains

To determine whether strains capable of hyphal growth differ from yeast-form counterparts in their interaction with the host, one representative fusion product from each cross was tested *in vitro* in the RHE infection model^10^, and *ex vivo* and *in vivo* with a murine model of *Malassezia* skin colonization^22^; the natural diploid hybrid CBS7019, and parental haploid strains were included as controls. In the RHE model, fusion products formed hyphae and invaded the cornified layer similar to CBS7019, whereas parental strains remained in the yeast form and were largely confined to the epidermal surface (**Fig. 6a**). SEM imaging of the RHE surface further reinforced this observation, revealing active tissue penetration and invasion by *M. furfur* hyphae (**Fig. 6b**). Consistent with invasive growth, barrier integrity assessed by transepithelial electrical resistance (TEER) and lucifer yellow (LY) permeability, was generally reduced in RHE infected with hypha-forming strains compared to the yeast-form parental strains, although significance was not reached for all strains tested in both assays (**Fig. 6c**). Hypha-forming strains also generally triggered higher expression of genes encoding proinflammatory cytokines IL-1β, IL-8, and IL-17C in RHE, as well as of genes encoding the antimicrobial peptides β-defensin 2 (β D2), β-defensin 3 (β D3), and LL-37, compared to the parental strains, although significance and magnitude varied across strains and markers (**Fig. 6d-e**).

**Fig. 6.**
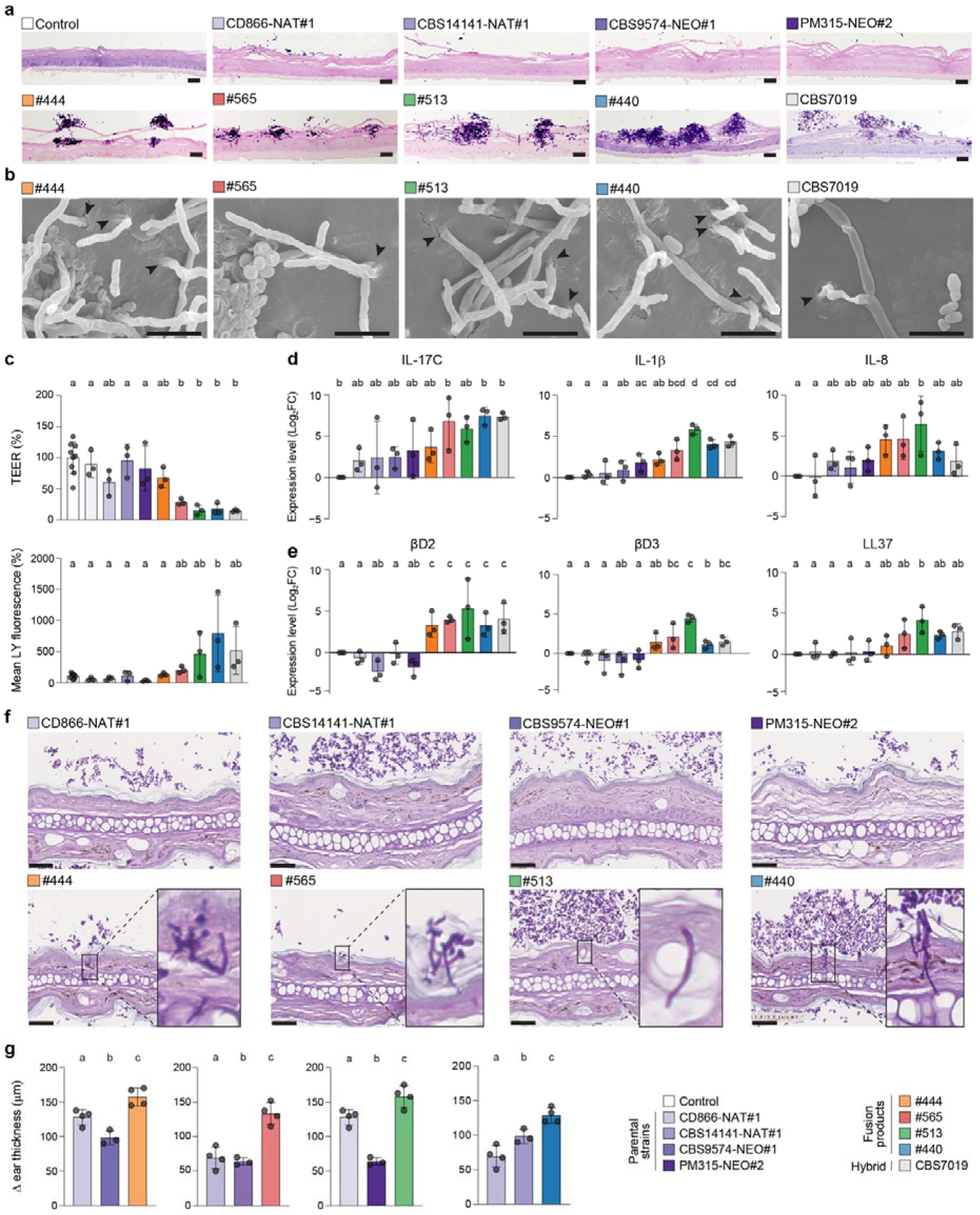
*M. furfur* hyphae-forming strains are more pathogenic than yeast-forming strains on human and mouse models. **(a)** RHE topically inoculated with *M. furfur* strains and stained with PAS staining on day 4 of infection. Scale bars, 50 μm. **(b)** Representative SEM images of hyphal strains during RHE infection; arrowheads indicate hyphae invading the cornified layer. **(c)** Barrier integrity assessed by transepithelial electrical resistance (TEER) and Lucifer Yellow (LY) permeability. Dots represents an individual measurement, and bars indicate mean ± SD. **(d, e)** Expression of genes encoding **(d)** proinflammatory cytokines (IL-1β, IL-17C, and IL-8) and **(e)** antimicrobial peptide (βD2, βD3 and LL37) in RHE infected with strains in the same order as in (a) and as given in the key. Expression levels (log_2_FC) were derived from Ct values relative to the endogenous reference *RPLP0* (ΔΔct method); bars show mean log_2_FC ± SEM, with dots indicating individual values from three biological replicates (*n* = 3). In panels c and d, statistical analysis was carried out using one-way Anova, with condition as the main factor and biological replicate as a blocking factor, followed by Tukey-adjusted pairwise comparisons; letters above bars indicate Tukey groupings (α = 0.05), with shared letters indicating no significant differences. Full pairwise comparison results are provided in Additional file 4. **(f)** PAS-stained histology sections of mouse ear skin colonized *ex vivo* with parental strains (top) or hyphae-induced fusion products (bottom). (**g**) Increase in ear thickness (Δ ear thickness, µm) of mice ear colonized *in vivo* with the same set of strains. Each graph pairs a fusion product with its parental strains. Dots represents an individual mouse; bars indicate mean ± SD. Statistical significance of the differences between groups was determined by ordinary one-way Anova followed by Tukey-adjusted pairwise comparisons; letters above bars indicate Tukey groupings (α = 0.05), with shared letters indicating no significant differences. Full pairwise comparison results are provided in Additional file 5.

The ability of *M. furfur* fusion products to penetrate skin tissue was next assessed *ex vivo* by infecting mouse ear skin with hyphae induced prior to inoculation, and by microscopic examination of periodic acid-Schiff (PAS)-stained histological sections. Parental strains in their yeast form were restricted to the stratum corneum surface, whereas fusion products hyphae extended through the epidermis (**Fig. 6f**), indicating that hyphal growth is associated with enhanced capacity for tissue penetration. *In vivo*, epicutaneous infection of mice ear with the same strains and measurement of ear thickness as a proxy for skin inflammation showed that hypha-forming strains consistently caused significantly greater ear swelling compared to their yeast-form parental strains (**Fig. 6g**). Taken together, these results demonstrate that *M. furfur* hypha-forming fusion products invade skin tissue more efficiently and elicit stronger host inflammatory responses than yeast-form parental strains, supporting a link between hyphal morphogenesis and pathogenicity during skin colonization.

## DISCUSSION

The skin fungus *M. furfur* has long been known to exist as both yeast and hyphal forms, with the latter associated with skin disease, yet the signals and genetic programs controlling this transition have remained poorly understood. Here, we show that skin-relevant lipids can trigger *MAT*-gene-dependent cell fusion and yeast-to-hyphae transition in *M. furfur*, and that intraspecific fusion products can undergo ploidy reduction and recombination reminiscent of a parasexual cycle, thereby linking host skin chemistry to fungal reproductive biology and pathogenesis.

Systematic dissection of individual RHE components identified olive oil as key trigger of hyphal formation in a natural diploid *M. furfur* strain. This effect is due to its abundance in unsaturated fatty acids, in particular oleic acid, which promotes *MAT* gene expression and the yeast-to-hyphae transition in strains with compatible *MAT* configurations within a single cell (**Fig. 1**). Other lipid sources, such as saturated fatty acids and Tween, commonly used to support *Malassezia* growth in axenic cultures, do not promote hyphal growth. Given that human sebum is rich in unsaturated fatty acids and squalene^23^, our findings suggest that the host-associated cutaneous lipids likely provide the environmental cue that triggers mating and hyphal growth, providing a mechanistic basis for understanding how naturally-occurring *M. furfur* hybrids form.

Consistent with this, skin-relevant lipids also promote mating between haploid *M. furfur* cells of opposite mating type. *M. furfur* comprises two parental lineages (P1 and P2) hypothesized to represent two distinct species, each with strains carrying either *MAT a1* or *a2* alleles but sharing a single *MAT B* allele (*b2* in P1; *b1* in P2)^18^. Applying a dual drug-resistance selection strategy, we show that strains from both lineages can undergo both intraspecific and interspecific mating. In both cases, the initial step of cell recognition, conjugation, and cell fusion between haploid isolates requires compatibility only at the *MAT A* locus **(Figs. 2, 7)**. Fused cells undergo karyogamy, generating a diploid monokaryon with a heterozygous genome, consistent with a chimera of homologous and homeologous parental genomes in intra- and interspecific fusion products, respectively. In fusion products that inherited compatible *MAT A* from both parents, a single *MAT B* allele, regardless of identity, is both necessary and sufficient for the yeast-to-hyphae transition **(Fig. 2, 7)**. The role of the *MAT A* pheromone/receptor (*P/R*) locus in cell recognition and fusion in *M. furfur* is hence consistent with its conserved role across basidiomycetes^12^; however, decoupling of *MAT B* from mating-type determination, while retaining its developmental role, represents a fundamental departure from canonical basidiomycete sexual development, and effectively renders *M. furfur* functionally bipolar.

In most basidiomycetes, hyphal development depends on heterodimerization of bE and bW homeodomain proteins derived from different *MAT B* alleles ^13,14^; the finding that a single allele suffices in *M. furfur* suggests that its HD regulatory mechanisms have been rewired to function independently of allelic complementation. While bE/bW self-recognition cannot be excluded, the consistently low expression of *bW* in different *M. furfur* strains under mating-inducing conditions (**Figs. 1, 5; Supplementary Fig. 11**) raises the intriguing possibility that a bE homodimer, rather than the canonical bE/bW heterodimer, drives hyphal development, as reported for bW/HD2 in the homothallic basidiomycete *Cystofilobasidium capitatum* ^24^. Notably, similar relaxation of *MAT B* allelic specificity has been observed in some *Microbotryum* anther-smut fungi ^25^, suggesting this may be a recurrent evolutionary strategy in basidiomycete fungi that have transitioned to non-canonical reproductive modes, or that have developed particular host-associations.

The genomic outcomes of cell fusion reveal a striking difference between interspecific and intraspecific crosses. Interspecific P1 × P2 fusion products are allodiploid hybrids with chromosome-wide heterozygosity and genome sizes of ∼16 Mb, resembling the naturally occurring *M. furfur* H2 lineage hybrids isolated from human skin ^18^, which suggests that H2 originated from interspecific mating events on the host. Interestingly, small LOH tracts were detected in fusion products from one of the two interspecific crosses (#440 and #503) but not the other, and affected different genomic regions in each strain, suggesting that LOH arises stochastically in *M. furfur* allodiploids, as previously observed in naturally occurring *M. furfur* hybrids ^18^. These events recall the interstitial (I-LOH) and terminal (T-LOH) LOH described in *S. cerevisiae*, arising from mitotic double-strand break repair, the former through gene conversion involving short exchanges and the latter resulting from crossing over or break-induced replication spanning large regions ^26^. Notably, the markedly lower levels of LOH in experimentally generated hybrids compared to natural *M. furfur* H2-hybrids suggest that LOH accumulates progressively after hybridization. A recent study on the H1-hybrid strain CBS1878 reveal that post-hybridization genome stabilization involves DNA repair-mediated chromosome restructuring coupled with homeologous recombination and gene conversion, generating localized LOH events ^27^. Therefore, LOH events are a dynamic driver of post-hybridization genome evolution, linking genome stabilization with the generation of genetic diversity that may facilitate adaptation in hybrid *M. furfur*. The experimental system established here now provides a unique opportunity to investigate the mechanisms and dynamics underlying these processes during the earliest stages of hybridization.

In contrast, fusion products of intraspecific crosses follow a different genomic trajectory: upon passaging without selection, diploid fusion products can undergo spontaneous ploidy reduction and recombination, generating haploid progeny with recombinant genomes. This may explain why diploids derived from intraspecific crosses have never been isolated from the skin. This process, however, is variable in both rate and genomic outcome: (i) some fusion products remained diploid even after passaging, (ii) others were haploids and with recombinant genomes, and (iii) others displayed mixed ploidy with progressive LOH accumulation, likely representing intermediate steps toward genomic stabilization (**Fig. 4**). Compared with the hybrids, fusion products obtained in intraspecific crosses exhibited a higher number of I-LOH and terminal T-LOH events that involved larger genomic regions, suggesting that their genomes are less stable and subjected to a different contribution of LOH-mediated stabilization. The similar LOH patterns observed across passaged isolates derived from the same fusion product are intriguing and could reflect several non-mutually exclusive processes. LOH events may have arisen very early after cell fusion, prior to single colony isolation, such that all passaged colonies inherited the same pre-existing LOH, although the largely heterozygous diploid genomes of the 0b samples suggest this would only have affected a minority of cells. Alternatively, the founder population (sample 0b) may already have been heterogeneous in ploidy and LOH status, with certain genotypes conferring a growth advantage on non-selective medium and being preferentially recovered as single colonies. Finally, specific genomic loci may represent hotspots for LOH, driven by selective pressure to eliminate deleterious alleles.

The generation of haploid progeny with recombinant genomes in intraspecific crosses is consistent with a parasexual cycle, in which cell fusion, nuclear fusion, and subsequent ploidy reduction generate genetically diverse haploid offspring without a canonical meiosis. Random segregation of *MAT* alleles in the haploid progeny likely ensures that both mating types are represented, enabling *MAT*-compatible offspring to re-enter the parasexual cycle and perpetuating genetic diversification (**Fig. 7**). Parasexuality in *M. furfur* shows notable analogies with the parasexual cycle of *Candida albicans*, in which, in the absence of canonical sexual structures, diploid cell fuse to form tetraploids that undergo ploidy reduction through concerted chromosome loss (CCL), a process in which chromosomes are lost in an apparently random and concerted fashion with recombination occurring between homologous parental chromosomes ^28,29^. Parasexual CCL in *C. albicans* is thought to involve chromosome non-disjunction during mitotic divisions, with the meiotic protein Spo11 generating DNA double-strand breaks that promote inter-homolog recombination in concert with Rec8 (a meiosis-specific cohesion subunit), together providing a mechanistic link between meiosis and parasexual chromosome loss ^30^. Interestingly, the *M. furfur* genome includes an ortholog of *SPO11* but lacks *REC8*, suggesting that Spo11 alone may drive parasexual ploidy reduction in *M. furfur*, potentially through a recombination mechanism that operates independently of meiotic cohesion. In *C. albicans*, parasex results in elevated recombination across all chromosomes, with very short recombination tracts ^31^, while in *M. furfur* not all chromosomes showed evidence of recombination in individual progeny (**Fig. 4**). Recombination was nonetheless detected on all chromosomes across the combined progeny of both intraspecific crosses, indicating that no chromosome is refractory to recombination per se. The lower per-chromosome recombination frequency in *M. furfur* compared to *C. albicans* may reflect biological differences between the two parasexual systems; it is also possible that, unlike canonical meiosis where at least one crossover per chromosome is typically required for proper segregation^32^, parasexual ploidy reduction in *M. furfur* does impose such a per-chromosome constraint, though this remains to be demonstrated. Lastly, the virulence data presented identify hyphal morphogenesis as a key determinant of *M. furfur* pathogenic interactions with the host, with hyphae-forming strains (regardless of whether they originate from intra or interspecific crosses) exhibiting enhanced tissue invasion, epidermal barrier disruption, and stronger immune activation compared to yeast-form counterparts in human and mouse skin models (**Fig. 6**). These findings are consistent with the known association of *M. furfur* hyphae with PV and seborrheic dermatitis^7,9^, and with a well-established link between morphological transitions and pathogenicity in fungal pathogens, such as *U. maydis* and *C. albicans* ^19,28^. However, haploid parasexual progeny did not form hyphae and are thus unlikely to contribute directly to acute tissue invasion. We speculate that parasex may instead provide an adaptive route in host-associated niches by generating genetic diversity without requiring the production of potentially immunogenic spores ^33^. Parasexuality and hyphal morphogenesis may therefore act as complementary processes: the former promoting population diversification over longer timescales, the latter driving tissue invasion and inflammatory host responses when skin-associated lipids induce *MAT*-associated hyphal development.

**Figure 7.**
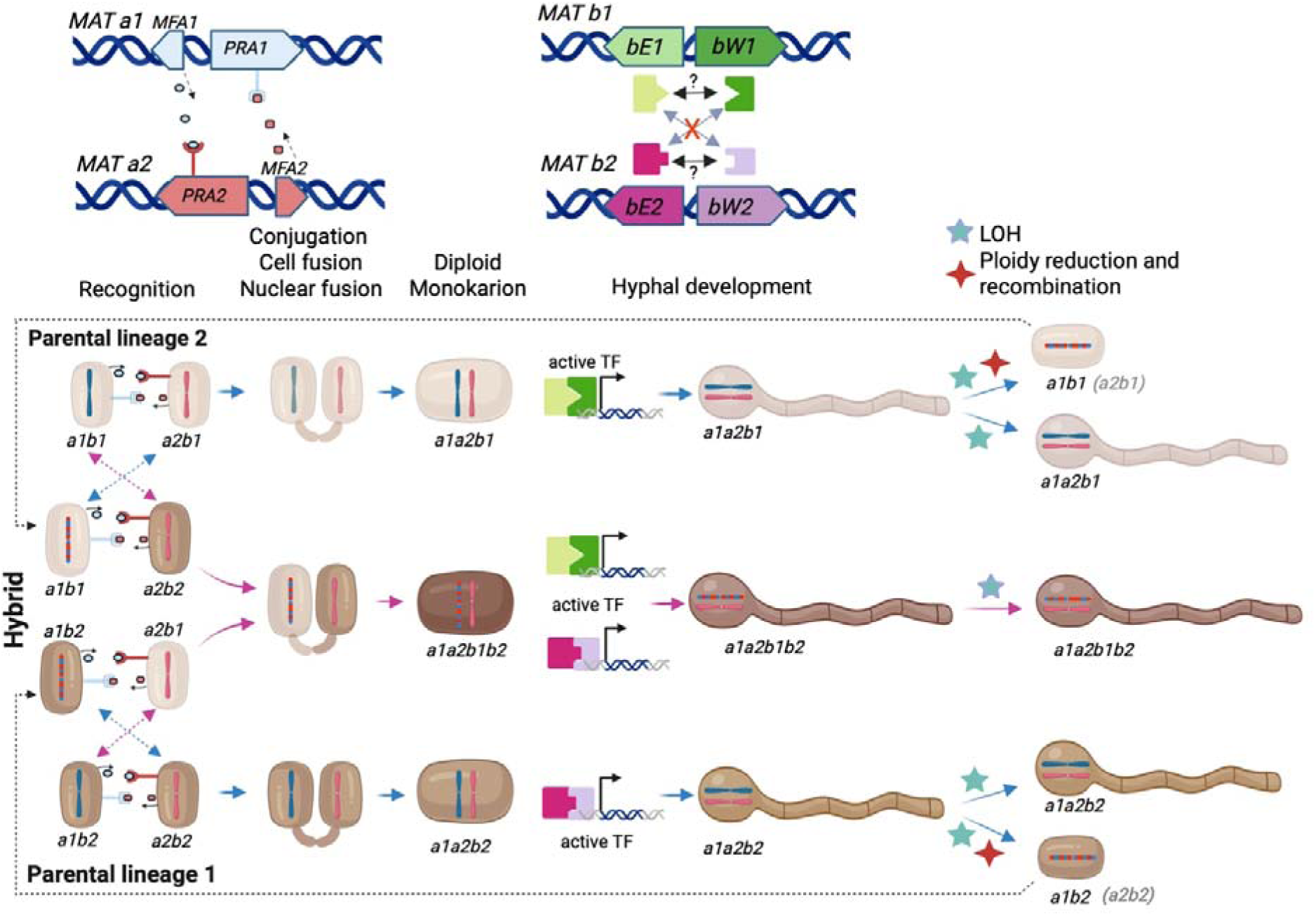
Model of *MAT*-dependent parasexual development in *Malassezia furfur*. The upper part of the figure summarizes the molecular machineries involved in *M. furfur* hybridization or parasexual reproduction and hyphal development, while the lower part shows the corresponding morphological changes and the *MAT* genotypes. Cell recognition, conjugation, and cell fusion are controlled by the *MAT A* locus through interactions between pheromone (encoded by *MFA* genes) and pheromone receptor (encoded by *PRA* genes) in *MAT-A* compatible partners. Following cell fusion and karyogamy, a diploid monokaryon is formed. These steps are in common in both inter and intraspecific crosses. Hyphal production is controlled by a single *MAT B* locus and does not require compatibility between *MAT B* alleles, hence excluding the bE/bW heterodimerization. Within each *M. furfur* parental lineages/species, intraspecific mating (blue arrows) between haploid cells of opposite *MAT A* alleles generates diploid monokaryotic cells carrying *MAT a1a2* genotype and the *MAT B* of the parentals (*MAT b1* for the parental lineage 2 and *MAT b2* for the parental lineage 1). These diploids undergo hyphal development and can subsequently experience loss of heterozygosity (LOH) while remaining heterozygous filamenting strains, or undergo LOH, ploidy reduction, and recombination, producing haploid parasexual progeny with recombinant genomes and a haploid *MAT* configuration. The parasexual products are mating compatible and can mate with *MAT A* compatible strains (dashed blue arrows), thereby completing the *M. furfur* parasexual cycle. Interspecific mating (indicated by fuchsia arrows) between strains belonging to the two parental lineages generates a hybrid diploid monokaryon. The model shows a mating between a parasexual product derived from parental lineage 2 (*MAT a1b1*) with a strain of the parental lineage 1 (*MAT a2b2*), generating a hybrid with a *MAT a1a2 b1b2* configuration. Hybrid cells undergo filamentation and accumulate LOH events during genome stabilization.

In conclusion, our findings point to a scenario in which the skin is not merely a passive substrate for *Malassezia* colonization, but rather an active arena for (para)sexual reproduction, hybridization, and genome diversification, initiated by the lipids that define this unique ecological niche, and adding growing evidence that host-associated fungi may rely on reproductive strategies distinct from canonical meiosis.

## MATERIALS AND METHODS

### Strains and culture conditions

*M. furfur* strains used in this study are listed in **Supplementary Table 1**. They were regularly maintained on modified Dixon’s media (mDixon) [mycological peptone (10 g/L), malt extract (36 g/L), glycerol (2 ml/L), tween 60 (10 ml/L), desiccated ox-bile (10 g/L), and agar (20 g/L) for solid media] with incubation temperature of 30°C.

*A. tumefaciens* strain EHA105 was used for *Agrobacterium*-mediated transformation of *M. furfur.* This strain was cultivated at 30°C in Luria Bertani [tryptone (10 g/L), yeast extract (5 g/L), NaCl (10 g/L), and agar (20 g/liter) for solid media], or LB supplemented with 50 µg/mL of kanamycin sulphate when transformed with binary vectors for *M. furfur* transformation.

### *Agrobacterium*-mediated transformation of *M. furfur*

*Agrobacterium*-mediated transformation of *M. furfur* was carried out as previously reported ^34^. Haploid *M. furfur* strain PM315 was transformed with the binary vector containing the *MAT a1* and MAT *b4* alleles together with a *NEO* marker ^16^. Haploid *M. furfur* strains CBS14141, CBS9574, CD866 and PM315 were transformed with *A. tumefaciens* containing the binary vectors pAIM2 and pAIM6 conferring resistance to nourseotricin (NAT) and neomycin G418 (NEO)^35^, respectively. To obtain mCherry-tagged cells of *M. furfur* strain CBS14141, transformation was carried out using EHA105 containing the bicistronic plasmid pJG201701 that confers resistance to NAT ^36^. In all cases *M. furfur* transformants were selected on mDixon supplemented with NAT [(100 µg/mL) clonNAT, AB-102L, Jena Bioscience] and NEO [(200 µg/mL) A6798, PanReac AppliChem].

*A. tumefaciens*-mediated targeted mutagenesis was carried out to mutate both the *MFA2* and *PRA2* genes of the *MAT A2* locus, and both the *bE1* and *bW1* genes of the *MAT B1* locus of *M. furfur* CBS14141, and of its derived self-filamenting strains GI156 and GI166; the *NAT* marker was used. Mutation of the *MAT B1* locus was also carried out in *M. furfur* strain PM315 using the *NEO* marker. Briefly, the 5ʹ and 3ʹ flanking regions for homologous recombination were amplified from the genomic DNA of *M. furfur* CBS14141, using primer pairs listed in **Supplementary Table 6**. The *NAT* and *NEO* cassettes were amplified, respectively, from plasmids pAIM2 and pAMI6 using primers ALID2078-ALID2081. The PCR products and the double-digested (*Kpn*I and *Bam*HI) binary vector pGI3 were transformed into *S*. *cerevisiae* FY834 using lithium acetate and PEG 3750, and correctly recombined plasmids identified by PCR were transformed by electroporation in *A. tumefaciens* EHA105 for transformation, as previously reported ^35^. *M. furfur* transformants obtained were colony purified on selective medium, subjected to phenol–chloroform–isoamyl alcohol (25:24:1) DNA extraction, and the correct replacement of the target loci was assessed by diagnostic PCR using primers reported in **Supplementary Table 6**.

### Conditions promoting hyphal growth and mating in *M. furfur*

To induce filamentous growth, cellular pellet of *M. furfur* strain CBS7019 from a mDixon agar plate was resuspended in sterile water, and 25 µL were spotted within a 50 µL-drop of olive oil (O1514, Sigma-Aldrich) placed on filamentation medium agar (MM) [Glucose (15 g/L), Ammonium sulphate (3.3 g/L), Magnesium sulfate (0.5 g/L), Potassium chloride (0.5 g/L), Potassium dihydrogen phosphate (1g/L), agar (20g/L)] and incubated at 30°C; hyphal formation was monitored daily using an optical microscope (Zeiss Axioscope 5 equipped with a Camera Axiocam 305 color, and with Illumination System Colibri 3 for fluorescence) with a 100 X objective lens (“N-Achroplan” 100x/1.25Oil M27) and with immersion oil.

To define how nutrients availability affects filamentation, CBS7019 was spotted within a 50 µL-drop of olive oil placed on 2% agarose- and 2% agar-based media in which all the MM components were tested alone or in combination; hyphal formation was monitored as described above after 3 days on incubation at 30°C.

To identify the lipid component(s) that triggers filamentation, CBS7019 cells were spotted on filamenting medium within a drop of different commercial oils (jojoba, peanut, corn, sunflower, colza, and coconut) bought in a local market, with the exception of stearic oil (Carl Roth). Subsequently, CBS7019 cells were spotted on filamenting medium [(with and without 1% of Tween 60) P1629, Sigma-Aldrich] and then few drops of oleic acid (7213.1, Carl Roth), linoleic acid (62230, Sigma-Aldrich), linolenic acid (6039.1, Carl Roth), or squalene (442785, Sigma-Aldrich) were applied on the yeast cells. Olive oil was always used as a positive control while the negative control was MM without olive oil; hyphal formation was monitored as described above after 3 days on incubation at 30°C.

Mating assays were carried out using the same procedure reported to induce filamentous growth. *M. furfur* strains resistant to NAT or NEO were resuspended in sterile water, their optical density adjusted to OD600 = 1 (about 1 x 10^9^ CFU/mL), mixed in equal amount in water, and 25 µL of the mix spotted within a 50 µL-drop of olive oil placed on filamenting medium agar as described above. At the defined incubation time, the cellular mix was collected, washed with water, and plated on mDixon supplemented with NAT (100 µg/mL) and NEO (200 µg/mL) to select fusion products. The single strain represented the negative controls. Quantification of cell fusion rate was carried out after 72 h of incubation, with five independent experiments (biological replicates) each consisting of two or three technical replicates. Data of each biological replicates was obtained by the means of the technical replicates, and subjected to ordinary one-way ANOVA with Tukey’s multiple comparison test.

Mating experiments were performed also on filamenting medium with few drops of oleic acid, linoleic acid, linolenic acid, or squalene applied on the yeast cells inoculated with a loop; olive oil was used as above (positive control). Three biological replicates were carried out. At the defined incubation time, the cellular mix was plated on double drug medium as reported above.

The percentage of hyphal production of two characterized fusion products for each cross was recoded after 3 days of incubation in the conditions reported above. Three or four independent microscopic observations were carried out for each strain. The number of total cells and total hyphae present in the observed sample were recorded and subjected to weighted one way ANOVA with Tukey’s multiple comparison test.

To observe a cell fusion event, the mCherry-tagged strain CBS14141 was crossed with *M. furfur* strains CBS9574 and PM315 according to the conditions reported above and subjected to microscopic observation after 24 – 72 h of incubation. Fluorescence microscopy was performed as reported above using a filter compatible with mCherry excitation (550/25 nm) and emission (605/70 nm). DIC and mCherry images were super-imposed using Adobe Photoshop.

### Scanning electron microscopy

Scanning electron microscopy was carried out as previously reported ^10^. The *M. furfur* strains were cultivated in filamenting medium with oil for 3 days or on RHE for 4 days (see below), and fixed in 2.5% glutaraldehyde and 0.1 M cacodylate buffer (pH 7.4) at 4C for 1 hour. Sample was rinsed three time for 10 min each in 0.2 M cacodylate buffer, and maintained in 0.2 M cacodylate buffer at 4°C for 48 h. Samples were gradually dehydrated in a graded ethanol series (30%, 50%, 70%, 80% and 100%), and dried overnight with hexamethyldisilazane. Gold-coated samples were then analyzed using 6010LV scanning electron microscope (JEOL).

### Nuclear staining of *M. furfur* yeasts and hyphae

Nuclear staining in *M. furfur* yeasts and hyphae was successfully achieved using a combination of the protocols reported by Lachke et al. ^37^ and Lin et al. ^38^, with some modifications. Briefly, cellular pellet of CBS7019 was placed on filamenting medium as reported above and incubated at least for 3 days at 30°C. Cells were collected and washed with sterile water to remove excess olive oil, then washed twice in HEPS balanced salt solution (HBSS: 10 mM HEPES, 150 mM NaCl), and incubated for 10 min in HBSS with 40% ethanol. Cells were centrifuged, the supernatant discarded, and the cellular pellet resuspended in 49 µL of 50% glycerol + 1 µL of a DAPI solution at final concentration of 1 µg/mL. 10 µL of treated cells were placed on a sterile microscopic slide, covered with a coverslip, and placed at 60°C for 40 min to aid penetration of DAPI solution into fungal hyphae. Fluorescence microscopy was performed as reported above using a standard DAPI filter. DIC and DAPI images were super-imposed using Adobe photoshop.

### PCR analyses

Fusion products obtained in the crosses using NAT and NEO resistant haploid strains, in unilateral crosses of CBS14141 *b1*Δ with CBS9574-NEO#2 and PM315-NEO#3, and in bilateral crosses CBS14141 *b1*Δ × PM315 *b1*Δ, were subjected to PCR to characterize the markers inherited following mating. Primers specific for the *MAT a1, a2, b1, b4* alleles were used according to the condition reported by Theelen et al. ^18^, while the *NAT* and *NEO* marker as reported by Ianiri et al. ^35^ (**Supplementary Table 6**). Primers specific for the *MAT* alleles were also used to further confirm the mutation of the target *MAT a2*Δ and *b1*Δ in strains CBS14141, GI156 and GI166.

To determine which parental mitogenome was inherited by fusion products obtained from interspecific and intraspecific crosses, primers targeting the *cox3–nad3* intergenic region and the *cob* exon2–exon3 intronic region were used according to the condition reported by Theelen et al. ^20^. Specific primers for *cox1* were designed to discriminate the mitogenome of strain CD866 from that of strain CBS9674 (**Supplementary Fig. 6**). PCRs were performed using EconoTaq PLUS 2x Master Mix (Lucigen) according to manufacturer instruction, with initial denaturation at 94°C for 2 min, denaturation at 94°C for 30 sec, annealing at 55°C for 30 sec, and extension at 72°C for 1 min/Kb.

### RT-qPCR and statistical analyses

Expression analysis of *MAT a1, a2, b1, b2* expression was carried out for several strains and conditions. *M. furfur* CBS7019 was collected from mDixon agar, resuspended in water and spotted on filamenting medium within a drop of olive oil, oleic acid, linoleic acid, linolenic acid, or squalene. Cells were incubated for 24 h and 48 h and then subjected to RNA extraction. RT-qPCR analyses of strains GI156 and GI166 and derived MAT *a2*Δ and *b1*Δ mutants, and of CBS14141 × PM315 and CBS14141 *b1*Δ × PM315 *b1*Δ, was performed by inoculating the strains with a loop within a drop of olive oil placed on filamenting medium with 24 h of incubation at 30°C. The comparative control condition was represented by filamenting medium with no oil inoculated either with the parental strains GI156 and GI166, or with the WT cross CBS14141 × PM315.

For RNA extraction, *M. furfur* cells were collected from plates and directly immerged in 1ml of TRIzol reagent (Invitrogen) according to the manufacturer’s instructions. To increase RNA yield, 200 µl of sterile glass beads were added to each tube and vortexed 2 times for 30 seconds before processing the samples. 500 µl of the aqueous phase was mixed with an equal volume of 70% ethanol. Total RNA was purified using the RNeasy Mini Kit (Qiagen). 500 ng of total RNA was retrotranscribed into cDNA using the SuperScript III Reverse Transcriptase kit (Invitrogen), and cDNA was 1:10 diluted in water. For keratinocytes, 1 µg of total RNA was retrotranscribed into cDNA using the SuperScript III Reverse Transcriptase kit, and cDNA was 1:20 diluted in water. Rt-qPCR were carried out using the Takyon No ROX SYBR Master Mix (Eurogentec), and with primers at a working concentration of 300 nM (**Supplementary Table 6**); 5 µl of cDNA were used for RT-qPCR. A “no-template control” and a water control were included for each target. Amplification protocol starts with 600s of pre-incubation at 95°C, followed by 45 cycles composed of 10 seconds at 95°C, 10 seconds at 60°C, and 10 seconds at 72°C. Technical and biological triplicates were performed for each sample. Gene expression levels were normalized using the reference genes *TUB2* for *M. furfur,* and *RPLP0* for keratinocytes, determined using the comparative ΔΔCt method, and expressed as log2 of the fold change.

For expression of *MAT a1, a2, b1, b2* in *M. furfur* CBS7019 grown with oil and individual fatty acids, statistical analysis was performed separately for each gene and time point on ΔCt values using an ANOVA model with condition as the main factor and biological replicate as a blocking factor, followed by Tukey-adjusted pairwise comparisons.

For expression of *MAT a1, a2, b1, b2* in GI156 and GI166, and in the bilateral crosses CBS14141 × PM315 and CBS14141 *b1*Δ × PM315 *b1*Δ, statistical analysis was performed separately for each gene on log_2_FC values using an ANOVA model followed by Sidak-adjusted pairwise comparisons.

### Reference genomes and construction of pseudo-references

To analyze the genotypes of *M. furfur* fusion products, genome assemblies from one of the parental strains served as references for read mapping and variant calling. High-quality chromosome-scale assemblies were available for two strains: CBS14141 and PM315 ^18^. These assemblies functioned as the references in comparisons where they corresponded to one of the parental strains. For the CD866 × CBS9574 fusion products, no high-quality genome assembly was initially available. To enable comparative analysis, a pseudo-reference genome for CD866 was generated by assembling Illumina reads de novo with SPAdes v3.15.3. The resulting scaffolds were then reordered using D-GENIES based on whole-genome alignment to CBS14141, ensuring a consistent chromosomal orientation for analysis. For reference or pseudo-reference assemblies other than *M. furfur* CBS14141, gene annotations were transferred from the CBS14141 reference genome using Liftoff v1.6.3 ^39^, which performs coordinate lift-over via whole-genome alignment with minimap2 v2.17 to support downstream visualization and gene-aware plotting.

### Mapping and Variant calling

Raw Illumina paired-end reads were trimmed with fastp v1.0.1 ^40^ to remove adapter sequences and low-quality bases. Reads shorter than 50 bp after trimming were discarded. Adapter sequences were automatically detected in paired-end mode. Quality-filtered reads were aligned to a selected reference or pseudo-reference genome of *M. furfur* with BWA-MEM v0.7.17, and the resulting alignments were sorted and indexed with Samtools v1.17. Variants were called independently for each sample using FreeBayes v1.3.10 ^41^, assuming diploidy. Sites were retained only if they met the following thresholds: minimum total depth ≥5, alternate allele count ≥2, base quality ≥20, and mapping quality ≥30. Resulting VCFs were further filtered with bcftools v1.17 to exclude variants with a quality score (QUAL) below 20.

### Parental marker identification and genotype classification

For each fusion product analyzed, we identified a set of high-confidence parent-informative SNPs by mapping sequencing reads from one parental strain (designated as parent 2) to the genome assembly of the other parental strain, which served as the reference (designated as parent 1) in the variant calling step. Variant calls for the parent 2 were filtered to retain only biallelic SNPs where the genotype was homozygous for the alternate allele (GT = 1/1), with a minimum quality score (QUAL) of 20, total read depth ≥5, and at least 2 supporting reads for the alternate allele. These SNPs represent positions where parent 2 differs from parent 1 and were used as informative markers for genotyping the fusion products. This filtering step was implemented in a custom Python script using pysam v0.22.0. Genotype calls at these parent-informative SNP positions were then extracted for each fusion product and classified into three categories: homozygous for parent 1, homozygous for parent 2, or heterozygous. For marker positions where no variant was detected in the fusion product, the genotype was inferred to be homozygous for parent 1, as the absence of a variant indicates identity with the reference allele.

### Genome-wide genotype visualization

Genotype classifications across the genome were visualized using custom Python scripts built with matplotlib v3.8.2, pandas v2.2.2, and Biopython v1.83. A matrix-style bar plot was generated to depict genotype states (homozygous parent 1, homozygous parent 2, or heterozygous) across each chromosome for all analyzed samples. To simplify the visualization of extended regions of consistent genotype, adjacent variants of the same class were optionally merged into blocks if separated by ≤5,000 bp. In addition to the genome-wide view, zoomed-in plots were also generated and plotted together with gene models from a GFF3 file to provide functional context.

### Coverage analysis and visualization

To quantify genome-wide variation in read depth across fusion products, we computed normalized read coverage in non-overlapping bins using bedtools v2.30.0. For each sample, read alignments (BAM files) were processed to calculate coverage in fixed-size genomic windows of 5,000 bp. Genomic bins were defined with bedtools makewindows, and per-bin coverage was computed with bedtools coverage. Coverage values were then normalized by dividing the number of reads in each bin by the sample’s genome-wide median read depth. This normalization step was implemented in a custom Python script using pandas v2.2.2 and Biopython v1.83. The resulting coverage profiles were visualized as heatmaps using matplotlib v3.8.2 and seaborn v0.13.2. Binned coverage values were clipped to a range of 0 to 2x relative to the median to reduce the influence of extreme values and enhance visual contrast. Final composite figures combining genotype classification and coverage heatmaps were arranged manually in Adobe Illustrator for clarity of presentation, using plots generated by the custom scripts described above.

### Fluorescence-activated cell sorting (FACS)

Ploidy was assessed by fluorescence-activated cell sorting (FACS) analysis as described by Theelen et al. minor modifications. *M. furfur* strains were grown for 4 days on mDixon agar at 30 °C; cells were collected in PBS, washed one time with PBS and fixed in 70 % ethanol at 4 °C overnight. Fixed cells were sonicated for 2 minutes and them washed with 1ml of NS buffer (10 mM Tris-HCl, pH 7.6, 250 mM sucrose, 1 mM EDTA, pH 8.0, 1 mM MgCl2, 0.1 mM CaCl2, 0.1 mM ZnCl2, 0.55 mM PMSF, 7 mM 2-mercaptoethanol) and then stained with propidium iodide (30 µg/mL) in 0.2 ml of NS buffer containing RNase A (1 mg/ml) at 4 °C overnight in the dark. A volume of 50 μl of stained cells were added to 0.5 mL of Tris-PI mix [482 µl 1M Tris pH 7.5 + 18 µl Propidium Iodide (1 mg/µl)]. The final cell suspension was filtered through Miracloth (Calbiochem # 475855) prior to analysis. Flow cytometry was performed using a BD FACSCanto II analyzer as previously described ^42^.

### *M. furfur* infection on reconstructed human epidermis

Reconstructed human epidermis (RHE) were obtained using human primary keratinocytes as previously described ^43^. RHE on day 11 of reconstruction were pre-treated with sterile olive oil prior infection by 10^4^ yeasts per RHE according to a previously developed protocol ^10^. Infected RHE were collected four days after infection. Infected and control RHE were collected in a 4% formaldehyde solution for at least 24h before processing. Periodic acid-Schiff (PAS) staining was used to highlight *Malassezia* cell wall and RHE were counterstained with hemalum as previously described ^44^. Barrier integrity of RHE during infection was assessed using trans-epithelial electrical resistance (TEER) and Lucifer Yellow (LY) assay as previously described ^45^. Keratinocyte RNA was extracted using TRIZol reagent (Invitrogen). RHE were cut from the cell culture insert and dissolved in 50 µl of TRIzol using a grinder (Nippon Genetics). 950ul of TRIzol was then added to reach a total volume of 1 ml. Samples were processed according to the manufacturer’s instructions. To ensure a pure and DNA-free RNA, 400 µl of the aqueous phase was collected and mixed with an equal volume of 70% ethanol. Total RNA was purified and subjected to DNAse treatment according to manufacturer’s instructions using the RNeasy Mini Kit (Qiagen). RNA concentration was determined by spectrophotometry using a Nanodrop device. RT-qPCR was carried out as described in the dedicated paragraph.

### *Ex vivo* and *in vivo M. furfur* infection on mice hear

Animal experiments in this study were conducted in strict accordance with the guidelines of the Swiss Animals Protection Law and were performed under the protocols approved by the Veterinary office of the Canton Zurich, Switzerland (license number ZH200/2024). All efforts were made to minimize suffering and ensure the highest ethical and humane standards according to the 3R principles ^46^. Female C57Bl/6 mice were mice purchased from Janvier Elevage and kept in individually ventilated cages under specific pathogen-free conditions at the Institute of Laboratory Animals Science (LASC, University of Zurich). *M. furfur* strains were grown on MM agar in 200 µl olive oil (2 × 10⁷ cells) at 30 °C for 7 days prior to infection experiments. The fungus was then collected from the agar and suspended homogenously. For *ex vivo* colonization assays, mouse ears were collected from euthanized mice, cut in half, and placed dorsal-side up in a 24-well plate. 50 µl fungal suspension was applied on top and the ears were incubated for 48 h at 30 °C under humid conditions. Samples were fixed overnight in 4% PBS-buffered paraformaldehyde, embedded in paraffin, sectioned sagittally (9 µm), and stained using periodic acid-Schiff (PAS). Slides were mounted using Pertex mounting medium (Biosystem) according to standard protocols. Histological images were acquired using a digital slide scanner (NanoZoomer 2.0-HT, Hamamatsu) and analyzed with NDP.view2 software. For *in vivo* infection experiments, epicutaneous colonization of mouse ear skin was performed as previously described ^22^. Mice were anesthetized by intraperitoneal injection of ketamine (65 mg/kg) and xylazine (13 mg/kg), and a 100 µl fungal suspension (corresponding to 1 × 10⁷ cells in olive oil) was applied epicutaneously to the dorsal side of each ear. Ear thickness was measured prior to colonization and then daily throughout the experiment using an Oditest S0247 0–5 mm measurement device (Kroeplin).

## ACKNOWLEDGMENTS

This work was supported in part by the LEO Foundation (grant # LF-OC-22_001060 to S.L.L. and G.I.), and by the grants NIH/NIAID R21 AI168672-02, R01 AI039115-28, and R01 AI050113-20 to J.H. J.H. is co-director and fellow of CIFAR program Fungal Kingdom: Threats & Opportunities.

