## Supplementary figures for "Host skin lipids trigger *MAT*-dependent mating, pathogenic hyphal growth, and parasexual reproduction of *Malassezia furfur*"

**Supplemental Figure legend**

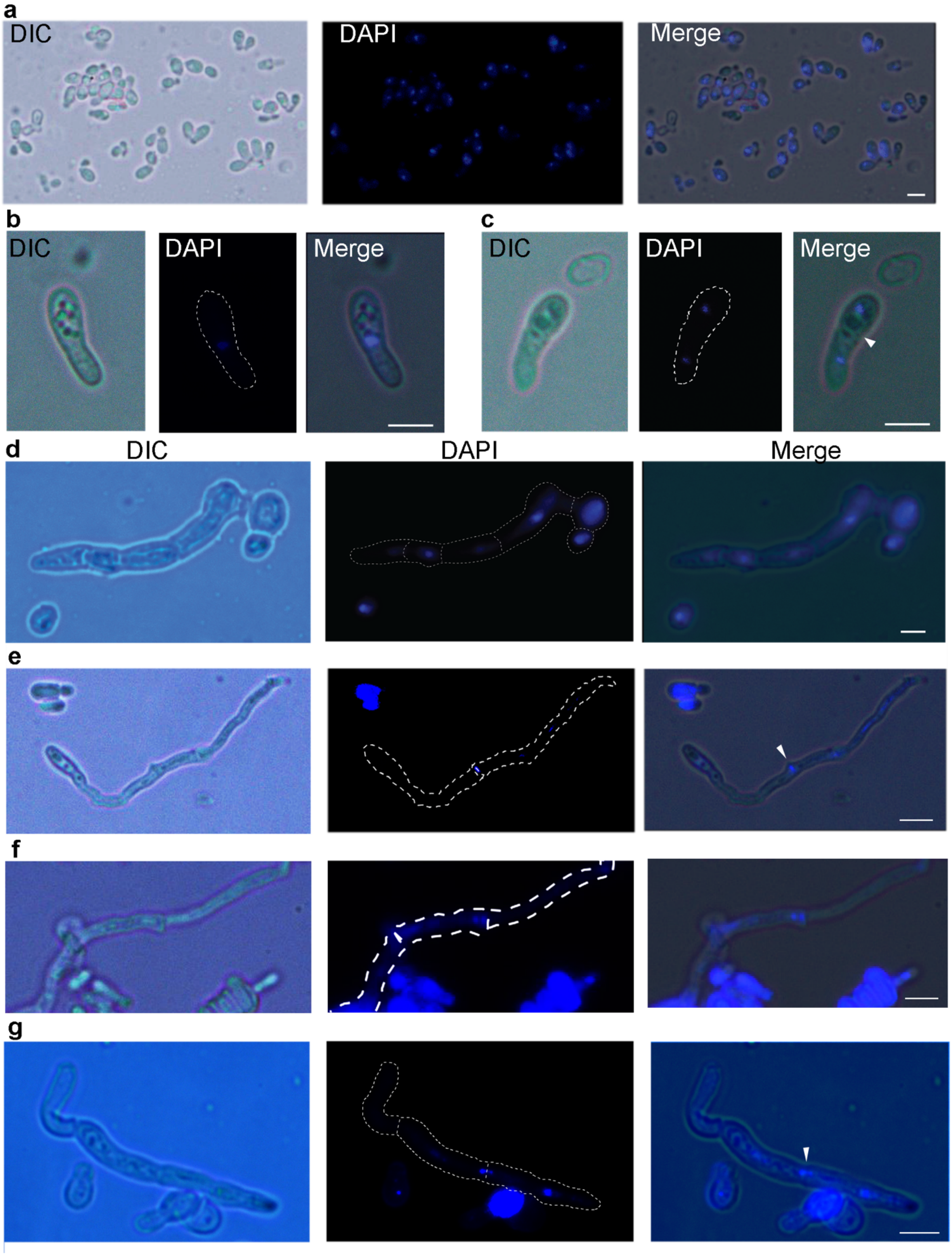


**Fig. S1: Nuclear dynamics in hyphae of *M. furfur* CBS7019.** All panels show differential interference contrast (DIC) light microscopy data (left), DAPI staining (center), and merged super-imposed images (right). **(a)** CBS7019 yeast cells grown under non-filamenting conditions. **(b)**

Germinating CBS7019 yeast cell showing nuclear migration toward the developing filament. **(c)** Germinating CBS7019 yeast cell after nuclear division, with one nucleus in the initial cell and one in the developing filament; the predicted septation site is indicated (arrowhead). **(d)** *M. furfur* hyphae with one nucleus in each cellular compartment. **(e)** *M. furfur* hyphae with one nucleus localized at the septum (arrowhead). **(f)** Portion of a *M. furfur* hyphae with a cellular compartment showing two nuclei close to a septum, suggesting that nuclear division takes place in one cellular compartment, and the novel nucleus migrates to the adjacent cellular compartment through the septum. **(g)** *M. furfur* hyphae showing two close nuclei, with one that crossed the septum (arrowhead). The bars indicate 5 μm.

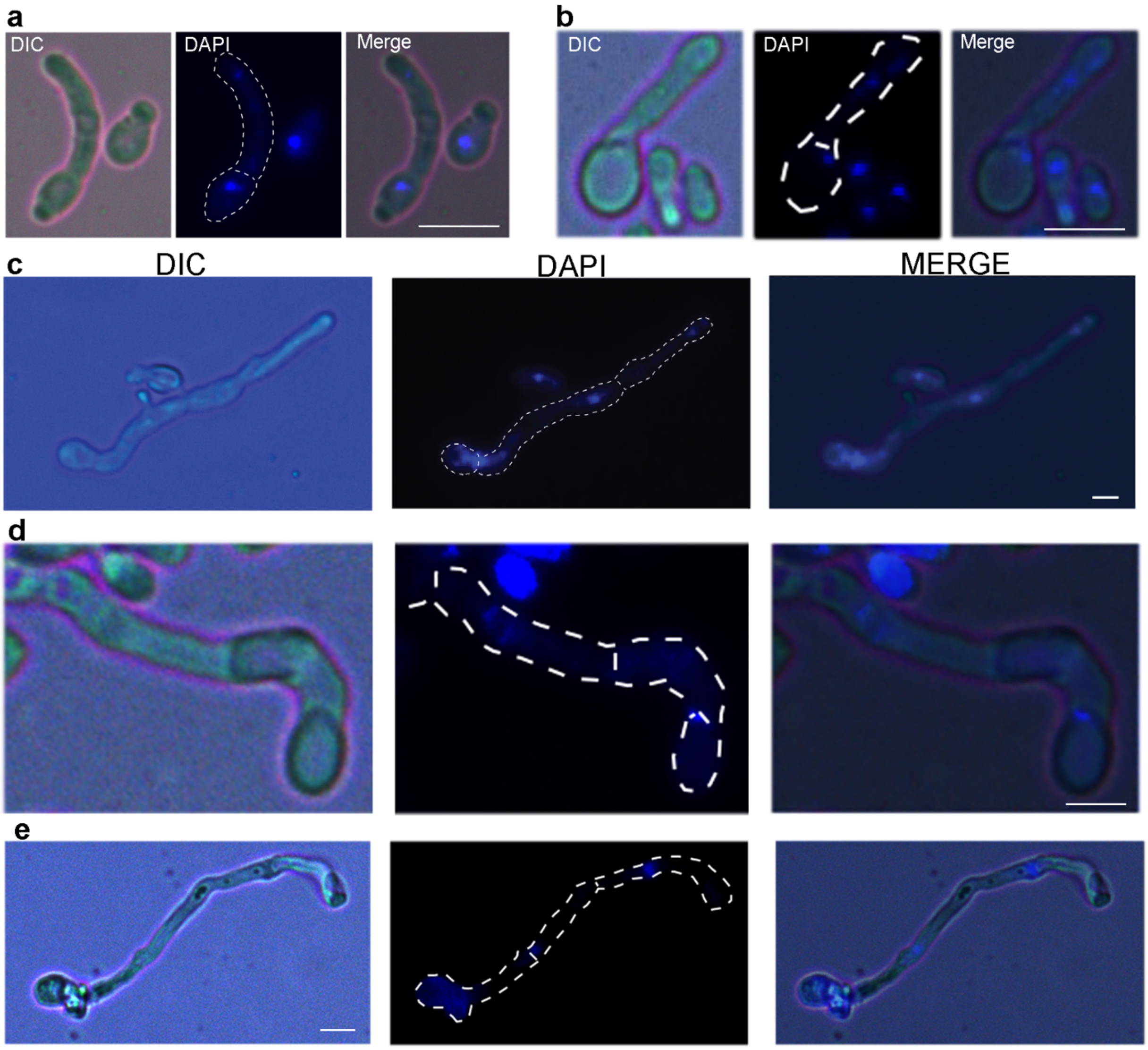


**Fig. S2: Nuclear dynamics in hyphae of the self-filamenting *M. furfur* strains GI156 and GI166.** All panels show differential interference contrast (DIC) light microscopy data (left), DAPI staining (center), and merged with super-imposition of DIC and DAPI fluorescence data (right). **(a)** *M. furfur* initial cell and hyphae of strain GI156 with each having one nucleus. **(b)** *M. furfur* strain GI156 initial cell with one nucleus, and hyphae with two nuclei, suggesting that a septum is formed after nuclear division. **(c-d)** *M. furfur* hyphae of strain GI156 with one nucleus at the septum between the initial cell and the first hyphal compartment; two nuclei are present in the last hyphal compartment, and a septum was being formed between them. **(e)** *M. furfur* hyphae of strain GI166 with one nucleus for cellular compartment; the nucleus in the first cellular compartment is not visible probably due to the presence of a yeast cell in front of the initial cell of the hyphae. The bars indicate 5 μm.

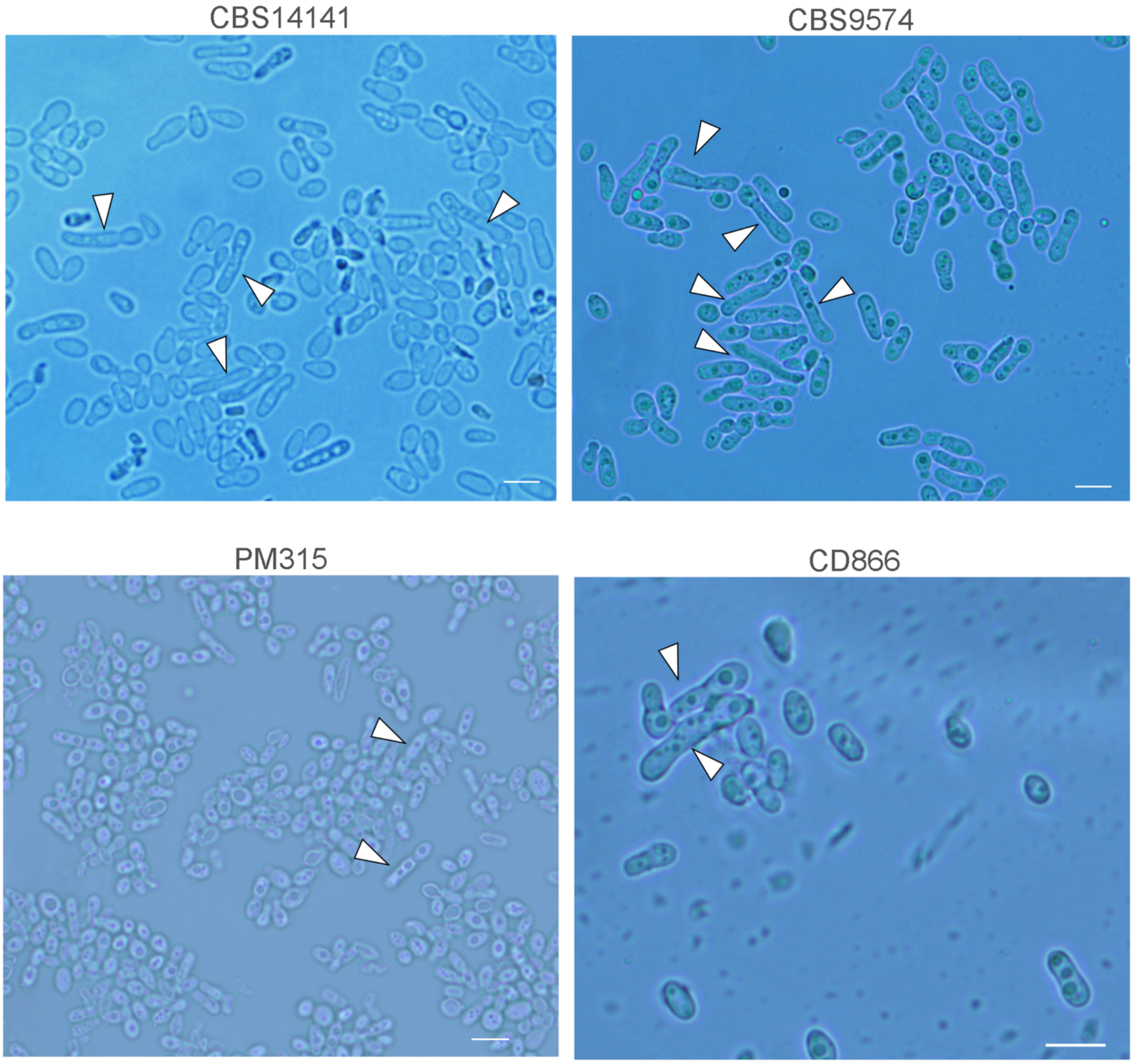


**Fig. S3: Formation of pseudohyphae in haploid strains of *M. furfur*.** Haploid *M. furfur* strains CBS14141, CBS9574, PM315, and CD866 were inoculated on filamentation medium plus olive oil without a mating partner and observed at the microscope after 3 days of incubation at 30ºC; arrowheads indicate pseudohyphae. Bars indicate 5 µm.

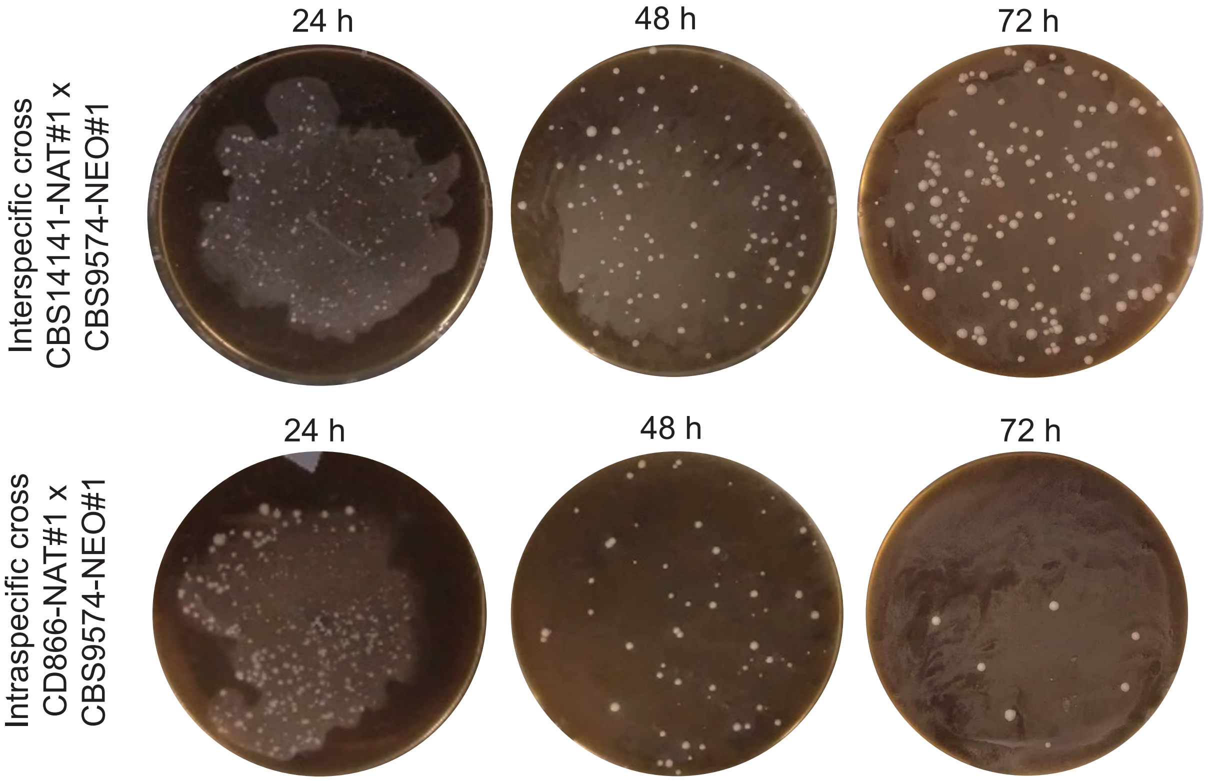


**Fig. S4: Cell fusion in *M. furfur* at different time intervals.** Selection of *M. furfur* fusion products obtained through interspecific CBS14141-NAT#1 x CBS9574-NEO#1 (P1 × P2) and intraspecific crosses CD866-NAT#1 x CBS9574-NEO#1 (P1 × P1) after 24, 48 and 72 h of incubation on filamentation medium plus olive oil.

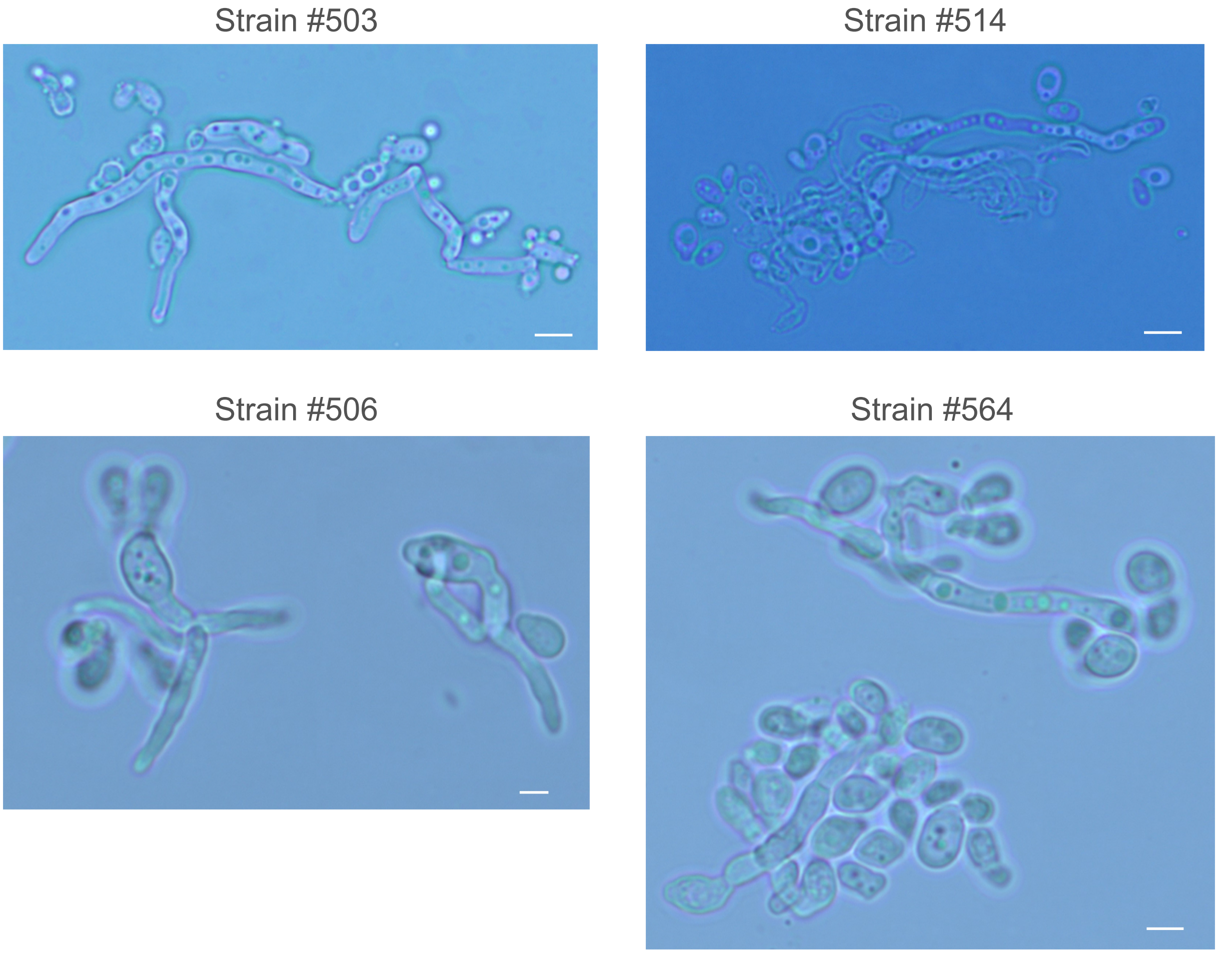


**Fig. S5: Hyphal morphology of *M. furfur* fusion products obtained through interspecific and intraspecific crosses.** Two fusion products obtained through interspecific (strains #503 and #514) and intraspecific (strains #506 and #564) crosses were inoculated on filamentation medium plus olive oil and observed at the microscope after 3 days of incubation at 30ºC; the bars indicate 5 µm.

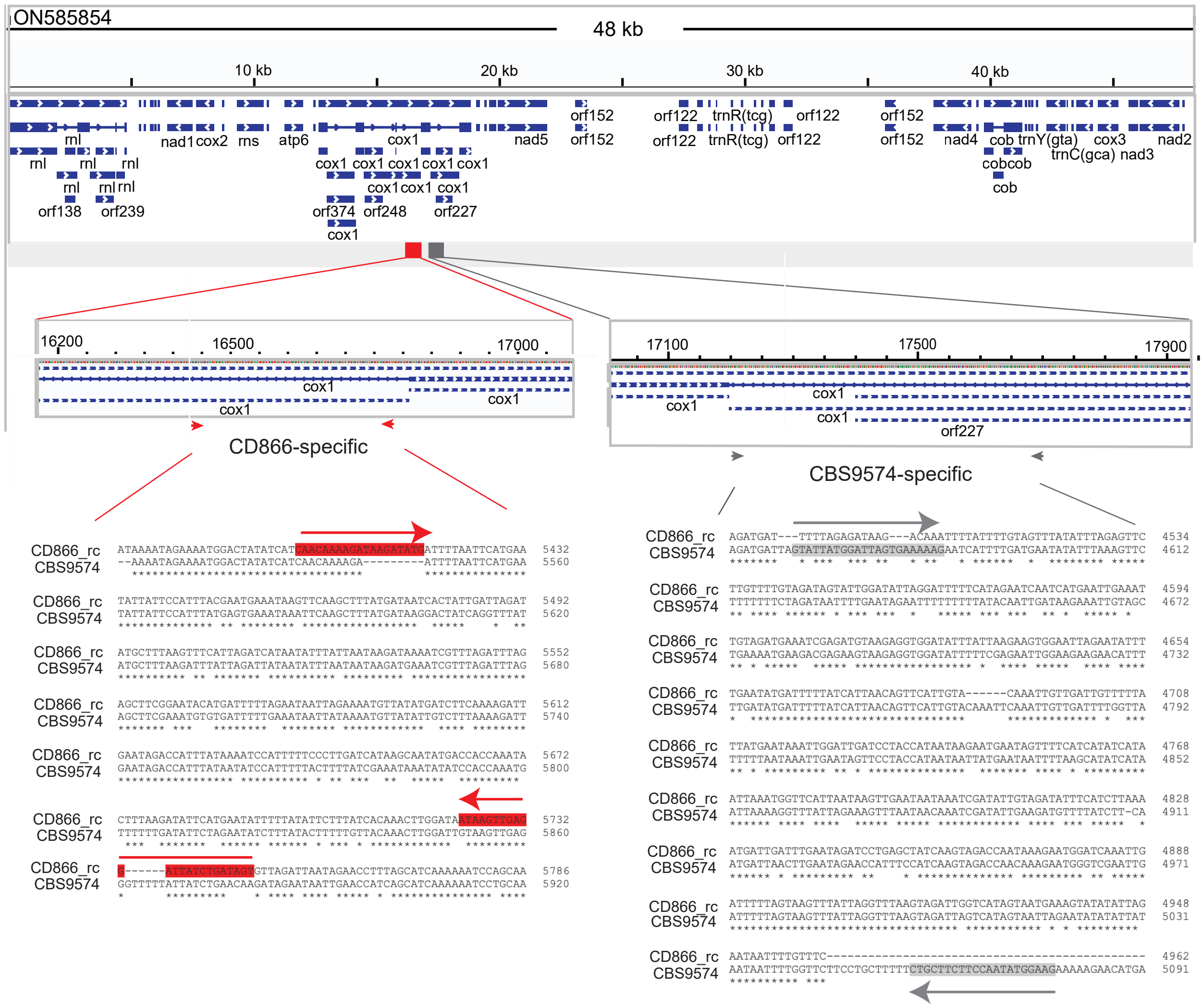


**Fig. S6: Development of a PCR assay to distinguish the mitochondrial genomes of the P1 *M. furfur* strains CD866 and CBS9574.** The mitogenome of *M. furfur* strains CD866 and CBS9574 was retrieved from de-novo assemblies of Illumina reads through comparison with the mitocondrial genome of *M. furfur* strain CBS9595 (accession ON585854). The retrieved mitochondrial sequences of strains CD866 and CBS9574 were aligned to identify divergent regions to design strain-specific primers that allowed to distinguish the two mitogenomes. For both strains the variability within the *COX1* gene was exploited. The figure shows the mitogenome annotation of *M. furfur* strain CBS9595 (accession ON585854), and the position of the primers specific for CD866 in red, and that of the primers specific for CBS9574 in grey.

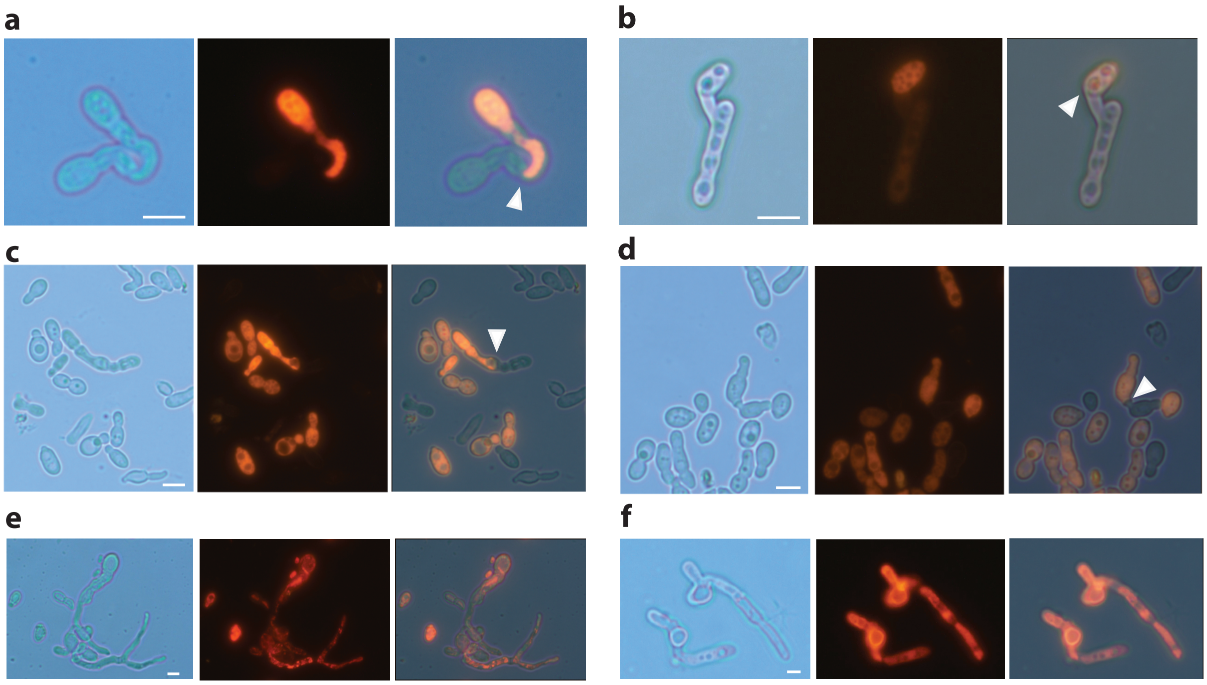


**Fig. S7. Cellular events involved in *M. furfur* cell fusion. (a – d)** The *M. furfur* strain CBS14141 was labeled with a mCherry fluorescent marker, crossed on filamentation medium with olive oil with non-fluorescent *M. furfur* strains CBS9547 (P2 × P1) and PM315 (P2 × P2), and observed at the microscope after 24 h of incubation. In **a – d** the arrowheads indicate contact points between the mCherry-tagged and the non-fluorescent *M. furfur* cells; all panels are relative to the cross CBS14141-mCherry × CBS9574, with the exception of panel b that reports the only connection found between CBS14141-mCherry and PM315. **(e - f)** Hyphae of the fusion products selected after crossing CBS14141-mCherry × CBS9574 **(e)** and PM315 **(f)**. All panels show DIC image (left), mCherry signal (middle), and merged super-imposed signal (right). Scale bars, 5 µm.

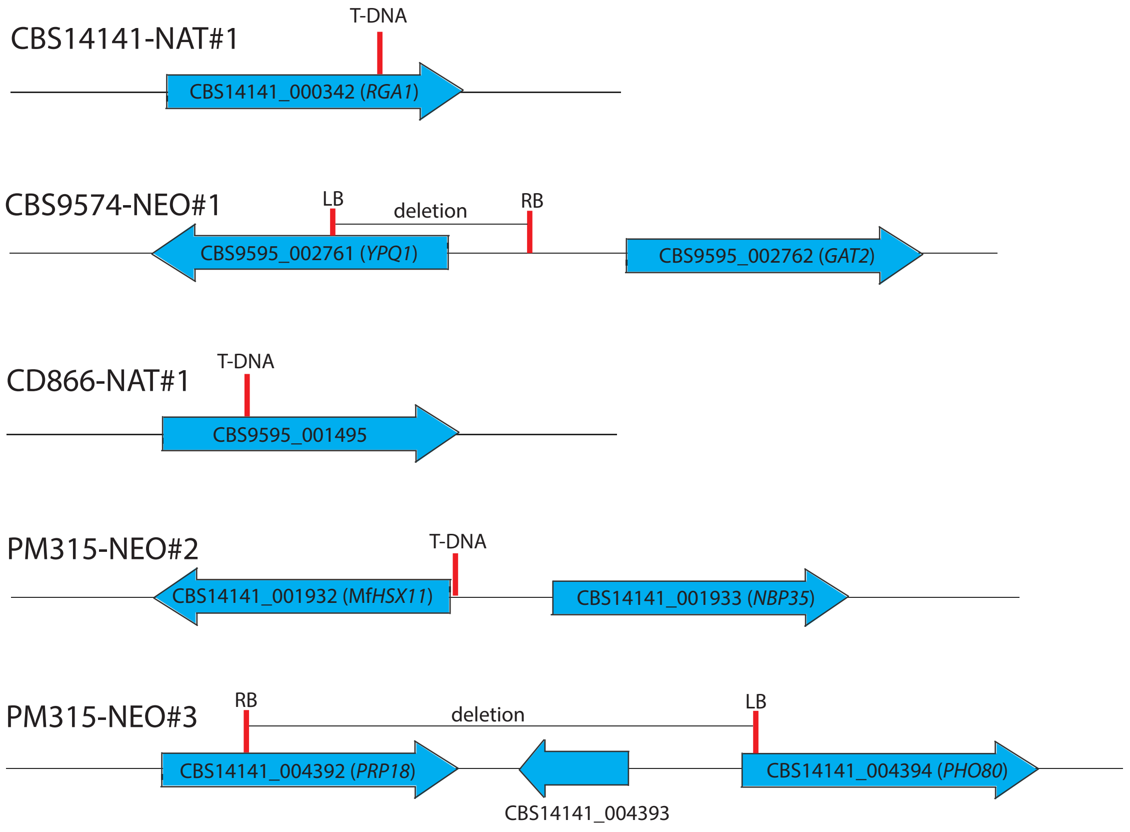


**Fig. S8: Identification of the mutated genes in the parental strains used in cell fusion experiments.** The genes bearing T-DNA insertions in the parental strains CD866-NAT#1, CBS14141-NAT#1, CBS9574-NEO#1, PM315-NEO#2, and PM315-NEO#3 were identified in the de novo assemblies of the Illumina reads through BLASTn analysis using the T-DNA as query. Retrieved sequences of P1 strains CD866 and CBS9574 were searched against the annotation of the reference P1 strain CBS9595, while retrieved sequences of P2 strains CBS14141 and PM315 were searched against the annotation of the reference P2 strain CBS14141. In all cases, the hit genes were used for BLASTp search in the *Saccharomyces* genome database to identify the closest ortholog and infer gene name, which is indicated in brackets where available. Where no *S. cerevisiae* orthologs were found, we reported the locus name when the gene encodes a hypothetical protein, or the gene name available on the *M. furfur* CBS14141 annotation, preceded by Mf (Mf*HSX11*).

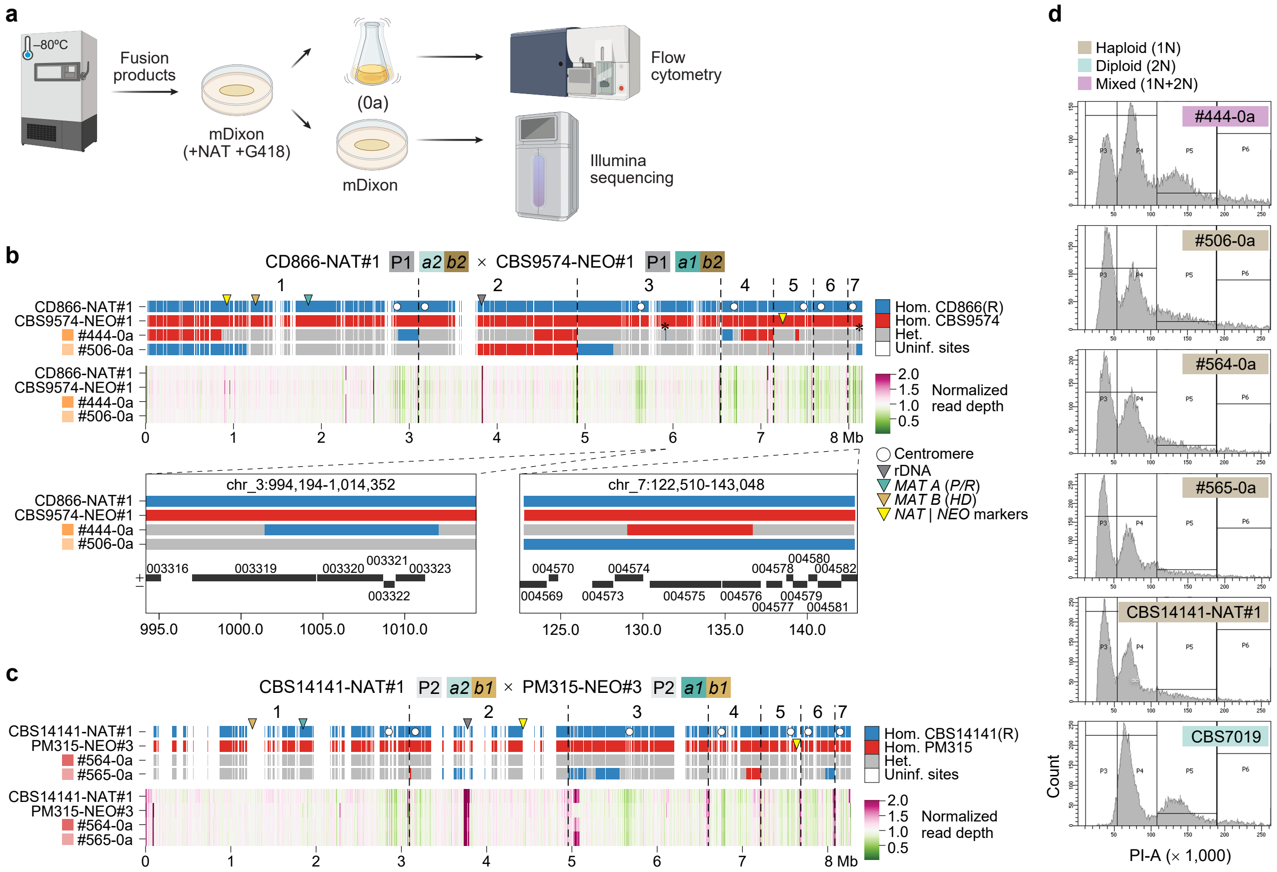


**Fig. S9: Initial genotype composition and normalized read coverage across *M. furfur* fusion products derived from crosses between marked strains within the same parental lineages (P1 × P1 or P2 × P2) that carry different *P/R* alleles but matching *HD* alleles. (a)** Experimental workflow used to characterize interspecific fusion products (labeled as 0a) through flow cytometry and Illumina Sequencing. Fusion products were revived from frozen stock (–80 °C) on mDixon plates supplemented with NAT and G418, then incubated in liquid mDixon for DNA extraction, and on solid mDixon for FACS. **(b)** Genotype classification (top tracks) and normalized read depth (bottom tracks) are shown for two independent fusion replicates of each intraspecific cross: **(b)** strains #444 and #506 obtained by crossing CBS9574-NEO#1 x CD866-NAT#1, and **(c)** strains #564 and #565 obtained by crossing CBS14141-NAT#1 × PM315-NEO#3. For each sample, genotype calls were classified as homozygous for the reference parent (blue), homozygous for the alternate parent (red), or heterozygous (gray), based on parent-informative SNPs. Adjacent variants of the same genotype class separated by ≤ 5 kb were merged into continuous segments. Regions identical between both parental strains are shown in white (uninformative sites). The reference genome used for each comparison is indicated with an "(R)". Read depth was calculated in non-overlapping 5 kb windows and normalized to the genome-wide median. Arrows indicate the reference-coordinate positions of the rDNA locus, T-DNA insertions, and *MAT A* and *MAT B* loci, as indicated in the key; T-DNA insertion sites were inferred from strain-specific *de novo* assemblies and projected onto the reference genome. White circles depict centromeres, and dashed lines indicate chromosome boundaries. Observed variability in coverage may reflect underlying differences in gene copy number or sequence divergence between parental genomes that affect read mapping. A magnified view in panel A shows a region of chromosome 3 and chromosome 7 exhibiting small localized loss of heterozygosity. Gene models shown in these zoomed regions were lifted from the *M. furfur* CBS14141 annotation and mapped onto the respective reference assemblies. All gene identifiers reflect the original CBS14141 locus_tag IDs. Strand orientation is indicated by "+" (forward/sense) or "–" (reverse/antisense) labels. **(d)** FACS analysis for ploidy determination in strains #506, #444, #564 and #565. The strains CBS14141-NAT#1 and CBS7019 were used as haploid and diploid control, respectively.

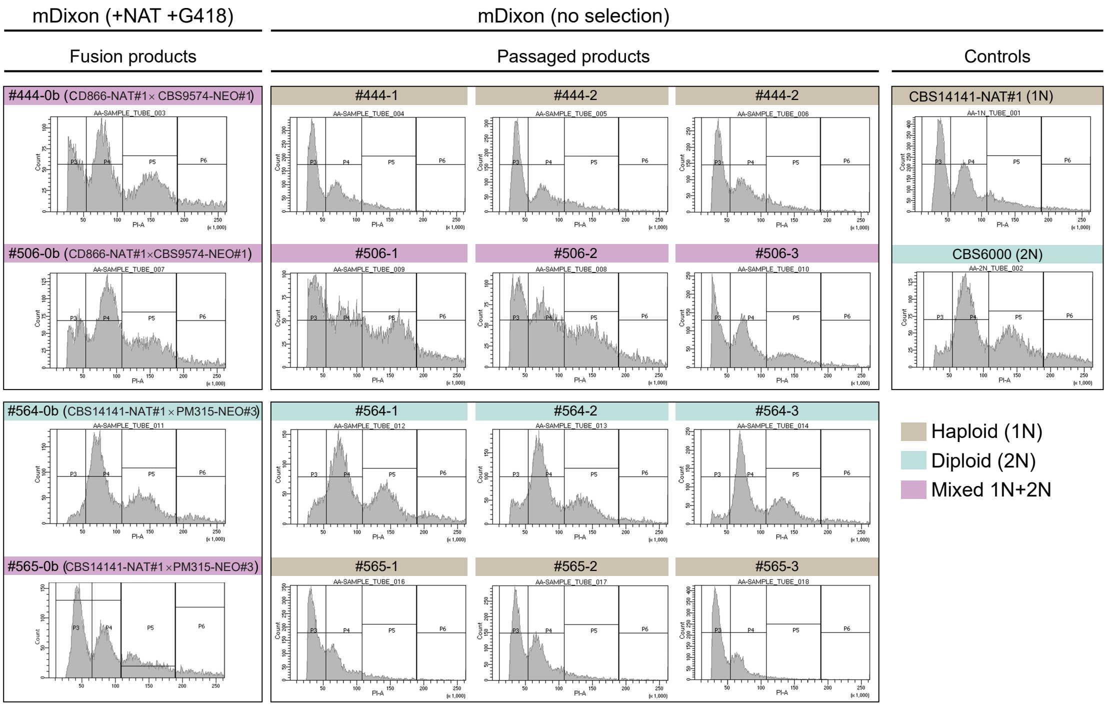


**Fig. S10: Flow cytometry for ploidy determination in the fusion products obtained through intraspecific crosses and in their passaged isolates.** The strains used for flow cytometry were obtained according to the representation reported in Figure 4a (strains 0b). Ploidy was determined through comparisons with the haploid and diploid controls *M. furfur* CBS14141-NAT#1 and CBS7019, respectively.

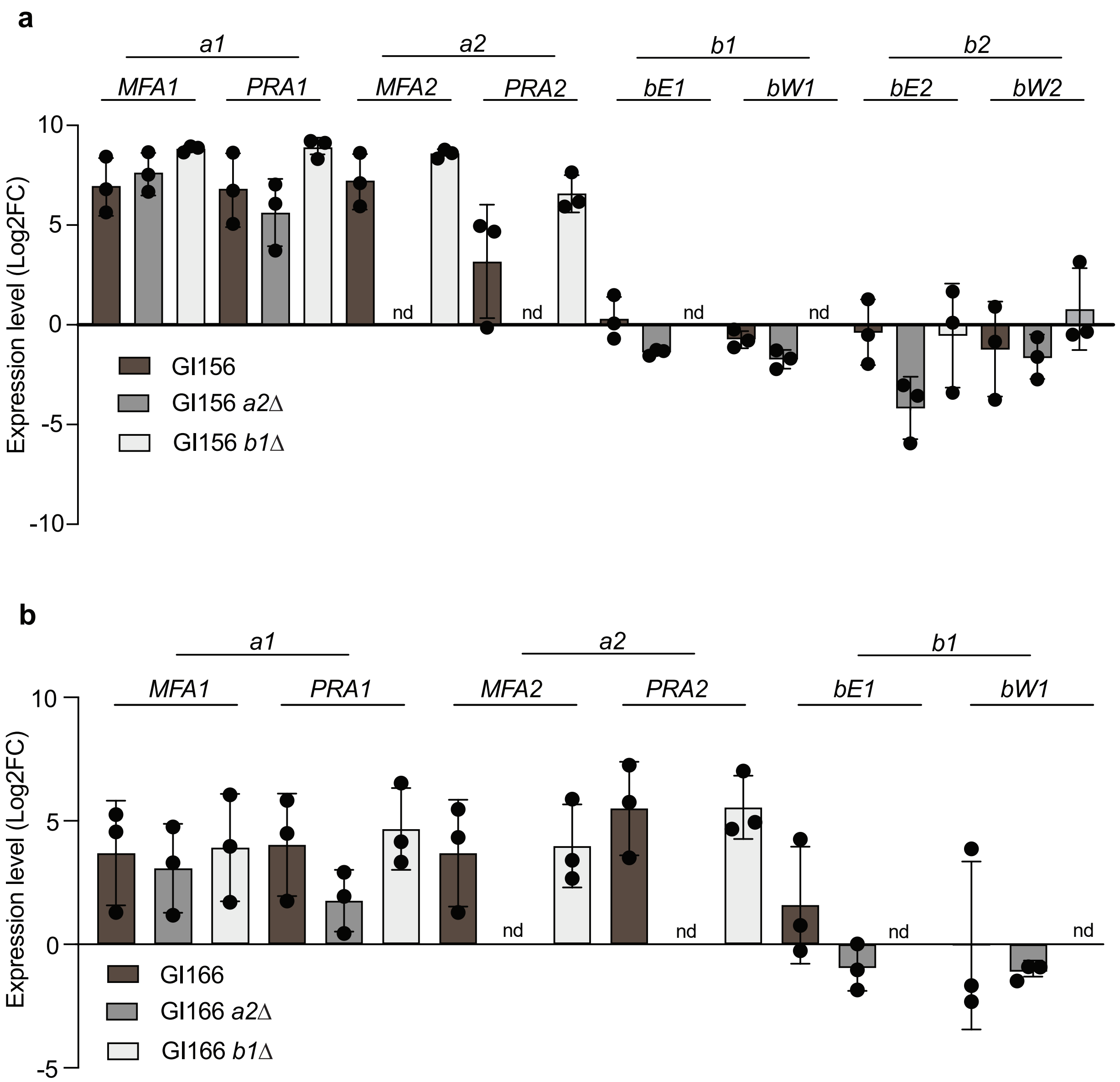


**Fig. S11: Expression levels of the *MAT* genes in strains GI156 and GI166, and their derived *mat a2*Δ and *mat b1*Δ mutants.** **(a)** RT-qPCR expression responses of the genes of the *MAT a1*, *a2*, *b1* and *b2* loci in strain GI156 and its derived *mat a2*Δ and *mat b1*Δ mutants after 24 h in filamentation medium alone (reference condition) or supplemented with olive oil. **(a)** RT-qPCR expression responses of the genes of the *MAT a1*, *a2*, and *b1* loci in strain GI166 and its derived *mat a2*Δ and *mat b1*Δ mutants after 24 h in filamentation medium alone (reference condition) or supplemented with olive oil. Ct values were normalized to the endogenous reference *TUB2* and converted to $log_2$ fold change ($Log_2FC$) relative to MM using the ΔΔCt method; because MM is the calibrator, its $log_2$ FC is defined as 0. Bars show the mean $Log_2FC$ across biological replicates ($n = 3$), with error bars indicating the standard error of the mean (SEM); dots indicate individual biological replicate values, each derived from the mean of three technical replicates. Statistical analysis was performed separately for each gene and time point on $Log_2FC$ expression values using an ANOVA model followed by Sidak-adjusted pairwise comparisons.
